# Translation is required for miRNA-dependent decay of endogenous transcripts

**DOI:** 10.1101/2020.01.21.913483

**Authors:** Adriano Biasini, Stefano de Pretis, Jennifer Y. Tan, Baroj Abdulkarim, Harry Wischnewski, Rene Dreos, Mattia Pelizzola, Constance Ciaudo, Ana Claudia Marques

## Abstract

Posttranscriptional repression by microRNA (miRNA) occurs through transcript destabilization or translation inhibition. Whereas RNA degradation explains most miRNA-dependent repression, transcript decay occurs co-translationally, raising questions regarding the requirement of target translation to miRNA-dependent transcript destabilization. To assess the contribution of translation to miRNA-mediated RNA destabilization, we decoupled these two molecular processes by dissecting the impact of miRNA loss of function on cytosolic long noncoding RNAs (lncRNAs). We show, that despite interacting with miRNA loaded RNA-induced silencing complex (miRISC), the steady state abundance and degradation rates of these endogenously expressed non-translated transcripts are minimally impacted by miRNA loss. To validate the requirement of translation for miRNA-dependent decay, we fused a miRISC bound lncRNA, whose levels are unaffected by miRNAs, to the 3’end of a protein-coding gene reporter and show that this results in its miRNA-dependent transcript destabilization. Furthermore, analysis of the few lncRNAs whose levels are regulated by miRNAs revealed these tend to associate with translating ribosomes and are likely misannotated micropeptides, further substantiating the necessity of target translation for miRNA-dependent transcript decay. Our analyses reveal the strict requirement of translation for miRNA-dependent transcript destabilization and demonstrate that the levels of coding and noncoding transcripts are differently affected by miRNAs.

## INTRODUCTION

Post-transcriptional regulation of gene expression by microRNAs (miRNAs) is widespread in eukaryotes and impacts diverse biological processes in health and disease [1, 2]. Most mature miRNAs are the product of a relatively complex biogenesis process. Primary miRNA transcripts, that generally depend on RNA Polymerase II for transcription, are initially processed by the nuclear enzyme DROSHA and its cofactor DGCR8 into a premature hairpin RNA of ∼60 nucleotides in length (pre-miRNA transcript) [3]. Pre-miRNAs are exported into the cytoplasm where they undergo a second round of processing by DICER resulting in a ∼22 nucleotide long double-stranded RNA duplex [4]. Loss of function mutations in any of the miRNA processing factors result in complete depletion of most miRNA species [5]. Argonaute proteins (AGO) bind mature miRNAs and guide target recognition of the RNA-inducing silencing complex (RISC). In mammals, target recognition relies primarily on complementarity between the miRNA seed region (position 2-8 of the mature miRNA) and miRNA recognition elements (MREs) in the target [6].

Posttranscriptional repression by miRNAs occurs by translation inhibition or transcript decay [2]. The contributions of RNA destabilization and translation inhibition to miRNA repression have been extensively studied [7, 8]. These studies support the general consensus that, translation inhibition precedes transcript deadenylation and decay [9–11], which in turn, is thought to account for most miRNA-dependent repression [9, 10, 12]. The coupling between translation inhibition and transcript destabilisation is further substantiated by evidence that protein-coding transcripts undergoing miRNA-dependent repression associate with translating ribosomes [13–19], and that most miRNAs loaded into RISC (miRISC) co-localize with polysomes [20–22].

These observations have raised questions regarding the requirement of translation for miRNA-dependent transcript decay. A number of experiments relying on the analysis of reporter constructs, revealed that transcript decay occurs even when translation initiation or elongation are impaired [23–25]. However, it is hard to reconcile the extent of target repression reported in these studies (up to five-fold) with the well-established impact of most miRNAs on endogenous transcript abundance, which rarely exceeds 2-fold [9, 10]. This has prompted concerns on whether exogenously expressed reporters faithfully recall the behaviour of most endogenously expressed transcripts.

To assess the requirement of translation for RNA destabilization of endogenous miRNA-targets and to overcome some of the limitations that may arise from using exogenous reporters, we took advantage of endogenously expressed cytosolic intergenic long noncoding RNAs, lncRNAs. This class of noncoding transcripts rarely associate with ribosomes [26] and have been previously shown to interact with miRISC machinery [27]. These transcripts thus provide a unique opportunity to address the outstanding question of whether miRNA-dependent decay occurs in the absence of translation. Specifically, we used 4-thio-uridine (4sU) to assess genome wide decay rates in wild-type (WT) and miRNA depleted cells. Our genome-wide analysis revealed that the decay rates of protein coding miRNA targets are significantly reduced upon miRNA loss whereas those of lncRNAs are only minimally impacted. Putative micropeptides were enriched among lncRNAs responsive to changes in miRNA abundance suggesting that translation is required for miRNA-dependent decay. We validated this hypothesis experimentally by inducing association of candidate lncRNA with translating ribosomes and found that this is sufficient to induce miRNA-dependent decay, further substantiating the prerequisite of translation for miRNA-dependent transcript decay.

## RESULTS

### Cytosolic lncRNAs interact with miRISC

Since posttranscriptional regulation by miRNAs occurs in the cytoplasm [6] and does not directly impact the levels of nuclear lncRNAs, we first classified lncRNAs based on their subcellular localization. We used RNA sequencing data from mESCs’ nuclear and cytosolic fractions [28] to estimate the expression of protein-coding transcripts (mRNAs) and intergenic long noncoding RNAs (lncRNAs) in these two subcellular compartments (Supplementary Figure S1A). We considered lncRNAs with a cytoplasmic/nuclear expression ratio higher than the median ratio for mRNA, which are predominantly located in the cytoplasm, to be cytosolic (n=1081). The remaining mESC lncRNAs, were considered to be nuclear (n=4953). Ribosome profiling data in mESCs [29] supports that mRNAs (50.4%) are more frequently associated with translating ribosomes than cytosolic or nuclear lncRNAs (6.6% and 4.0%, respectively, two-tailed Chi-square test, p-value< 10^-4^, Figure 1A). We took advantage of publicly available AGO2-CLIP [30] data for wild-type and DICER knockout mESCs, to assess whether cytosolic lncRNAs are associated with miRISC. We found that the fraction of mESC expressed cytosolic lncRNAs and mRNAs with experimental evidence for AGO2 binding is similar, (6% and 7% respectively, two-tailed Chi-square test, p-value=0.16), as is the density of bound sites within cytosolic lncRNAs (1.0 sites per kb of sequence) and mRNA 3’UTRs (0.7 sites per kb of sequence, two-tailed Mann-Whitney test, p-value<0.05, Figure 1B). Our ability to detect binding by miRISC, using this approach, is in part limited by the endogenous expression of transcripts as highlighted by the significantly higher expression of transcripts bound by AGO2 (average expression (TPM) bound=9.0 vs unbound=5.4, two-tailed Mann-Whitney test, p<2X10^-26^ Supplementary Figure S1B). Since lncRNAs are in general more lowly expressed than mRNAs, the proportion of lncRNAs bound by AGO2 may be higher than what is detected. The fraction of cytosolic lncRNAs bound by AGO2 with (6%) and without (7%) experimental evidence of ribosomal association is statistically indistinguishable (two-tailed Fisher’s exact test p=0.8), suggesting that AGO2 binding is independent of translation. We conclude, that consistent with previous analysis, most cytosolic lncRNAs do not stably associate with translating ribosomes [26], but are nevertheless targeted by miRISC [27], and are therefore, uniquely suitable to assess the impact of miRNAs on endogenous transcript destabilization in absence of translation.

**Figure 1.**
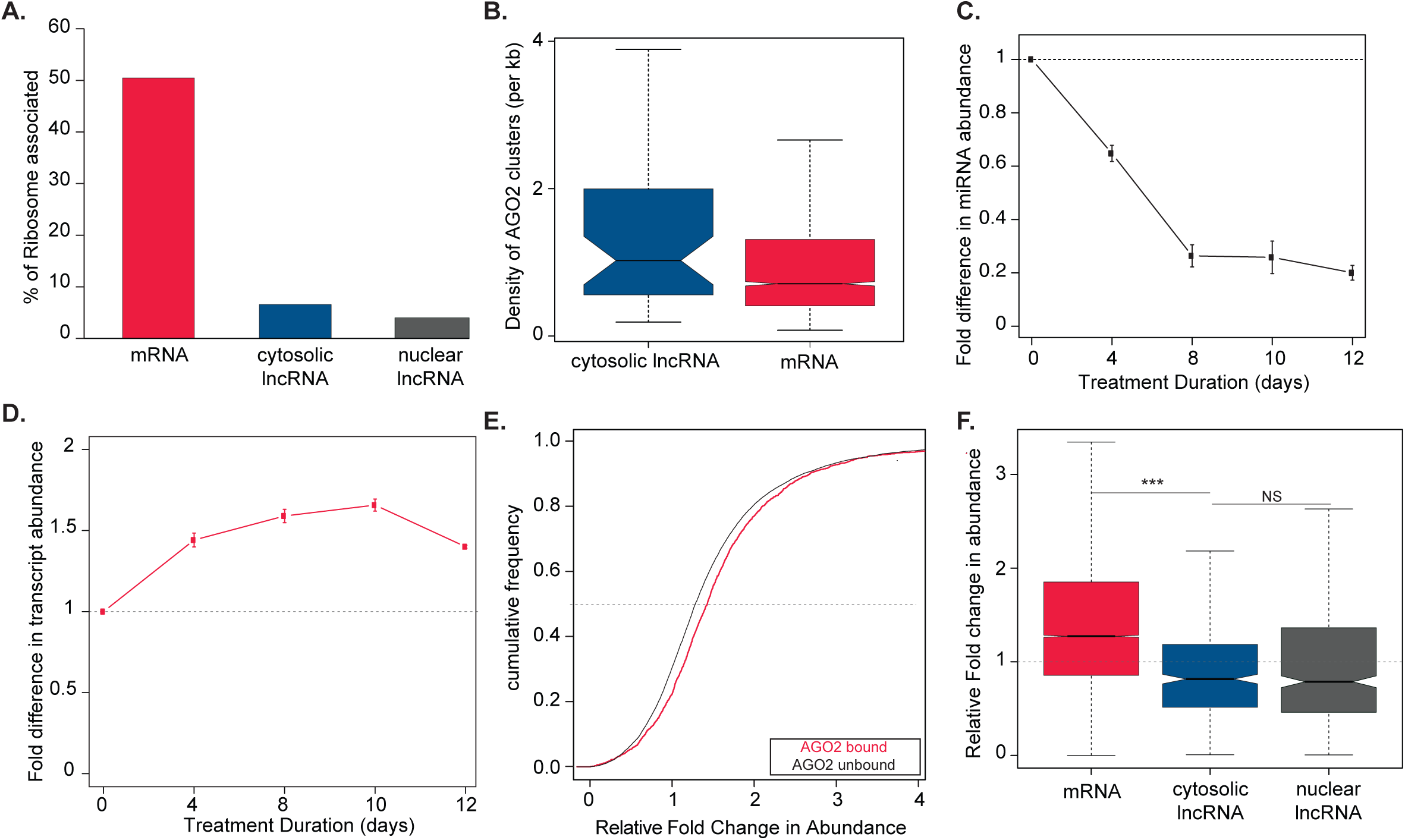
Steady-state abundance of lncRNAs is not directly affected by miRNA loss. (A) Percentage of mRNAs (n=6701, red) and predominantly cytosolic (n=57, blue) and nuclear lncRNAs (n=175, grey) with experimental evidence of binding by ribosomes (Translation Efficiency>0) in mESCs. (B) Density of AGO2 wild-type specific clusters across cytosolic lncRNAs (n=48, blue) and the 3’untranslated regions of mRNAs (n=2355, red) with experimental evidence for AGO2 binding in mESC (>0 AGO2 clusters). Small RNA and Poly(A)-selected RNA sequencing based estimates of the fold difference (y-axis) in (C) miRNA and (D) mRNA expression, respectively, relative to day 0, during a 12 days’ time-course (x-axis) following treatment of DTCM23/49XY mESC with 4-OHT and loss of DICER function. Points represent the average miRNA or mRNA expression and error bars the standard deviation based on 3 independent biological replicates. (E) Cumulative distribution plot of the fold-difference in expression after 8 days of tamoxifen treatment for mRNAs, expressed at day 0 (tpm≥1) with (n=1612) and without (n=12301) AGO2 clusters. (F) Distribution of the relative fold-change after 8 days of 4-OHT treatment in steady state abundance, relative to day 0, for mESC expressed (tpm≥1) mRNAs (n=19306, red), cytosolic (n=445, blue) and nuclear (n=529, grey) lncRNAs. Statistics: *-p<0.05, **-p<0.01 and ***-p<0.001.

### Steady-state expression of noncoding transcripts is minimally impacted by miRNAs

We first sought to determine whether cytosolic lncRNA expression was post-transcriptionally regulated by miRNAs. We took advantage of a mESC cell line containing two Cre/LoxP sites flanking the *Dicer* RNAse III domain on exon 21, and a *Cre* recombinase gene expressed under the control of a 4-hydroxytamoxifen(4-OHT)-inducible promoter [31, 32]. Exposure of these cells to 4-OHT leads to LoxP site recombination and strong depletion of DICER (Supplementary Figure S1C). Conditional loss of DICER function minimally impacts cell proliferation (Supplementary Figure S1D) and the transcript and protein levels of (Supplementary Figure S1E-G) of the pluripotency transcription factors, Nanog, Oct4 and Sox2. In contrast to what was previously reported for *Dicer* constitutive knockdown mESCs, that exhibit an 10-fold downregulation of *c-Myc* expression [33], in conditional *Dicer* mESC mutants the expression of this gene is only minimally impacted (Fold-change between KO and WT > 0.5, Supplementary Figure S1E), supporting that this system is better suited to investigate the direct effects of miRNA depletion.

We profiled small RNA expression following DICER loss of function and found that 8 days after 4-OHT addition, mature miRNA levels are reduced by ∼80% (Figure 1C). We validated these results, by RT-qPCR, for miR-290 and miR-295, which are among the most abundant miRNAs in mESCs [34] (Supplementary Figure S1H). Decreased levels of these miRNAs is associated, as expected, with a significant increase in the levels of some of their well-established targets [35] (Supplementary Figure S1I).

To assess the genome-wide impact of miRNA loss on mRNA and lncRNA expression, we used data from our previously published transcriptome-wide expression profiling following loss of DICER experiment in these cells [28]. As expected, and consistent with the role of miRNAs on posttranscriptional repression of protein-coding gene expression, we found that mRNA levels increased moderately but significantly following *Dicer* loss of function (Figure 1D). The fold-increase in expression, relative to control, in miRNA depleted mESCs is significantly higher (two-tailed Mann-Whitney test, *p*<1.4X10^-10^) for transcripts with experimental evidence for AGO2 binding (Figure 1E), supporting that the observed changes in mRNA expression are, at least in part, a consequence of mRNA alleviation from miRNA-mediated repression. In contrast to mRNAs, we found that lncRNA expression was minimally impacted by miRNA depletion (Figure 1F). Specifically, and in contrast to mRNAs, lncRNA steady-state abundance is slightly decreased in miRNA depleted cells (Figure 1F). This small decrease is likely an indirect effect of miRNA loss. Specifically, decreased levels of miRNAs are expected to result in increased steady state abundance of targets as observed for mRNAs (Figure 1F), whereas the impact of miRNA depletion is similar for both subcellular classes of lncRNA independent of co-localization with miRISC (Figure 1F). We conclude that, despite interacting with miRISC, cytosolic lncRNA transcript levels are not directly controlled by miRNAs (Figure 1F).

### No evidence for miRNA-dependent destabilization of noncoding transcripts

Steady-state transcript abundance depends on the rates of transcription, processing and degradation but only the degradation is directly controlled by miRNAs. To determine transcriptome-wide differences in degradation rate between miRNA depleted and control mESCs we performed, in duplicate, 4-thio-uridine (4sU, 200uM) metabolic labelling of RNA for 10 and 15 minutes, on mESCs 8 days after induction of DICER loss of function and control mESC. We sequenced total RNA and quantified intron and exon expression transcriptome wide from the pre-existing and newly synthetized RNA fractions. (Figure 2A, Methods). Principal component analysis of the gene expression estimates across the different samples revealed that the RNA fraction is the strongest discriminator between estimates followed by miRNA content and lastly by biological replicate (Supplementary Figures S2A-C). Degradation rates, that we estimated using INSPEcT ([36], methods) for the two different pulses durations (10 and 15 minutes) are highly correlated for both cell types (R^2^>0.75, Figure 2B-C). We used an alternative method (transcription block by Actinomycin-D) to validate the estimated differences in transcript stability between wild-type and miRNA depleted cells for a subset of transcripts spanning a range of fold-differences in degradation rates (Pearson R^2^=0.58, Supplementary Figure 2D).

**Figure 2.**
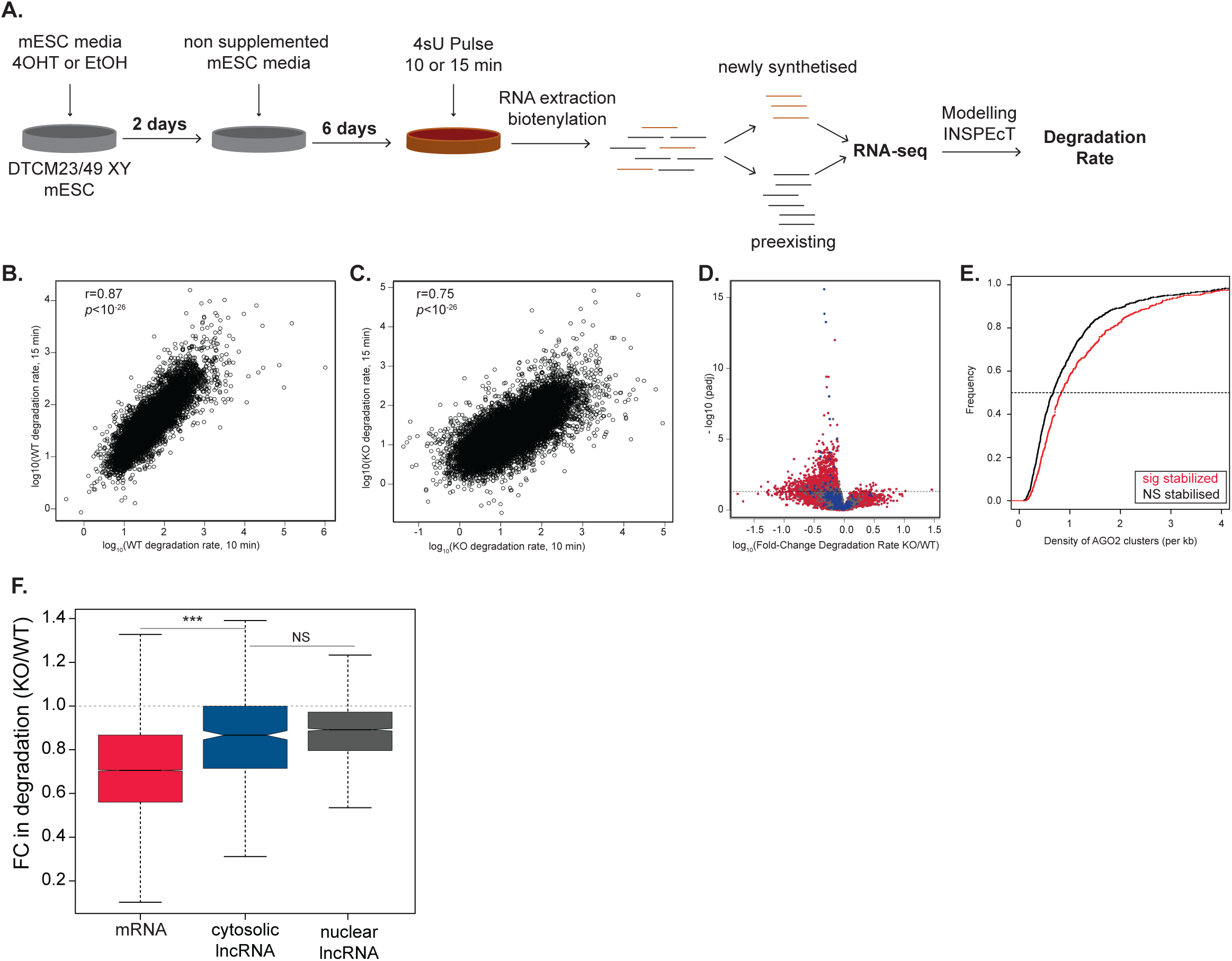
No evidence for miRNA-dependent destabilization of cytosolic lncRNAs. (A) Schematics of 4sU metabolic labelling of conditional *Dicer* knockout and wildtype cells experiment. Correlation between degradation rates (log10) obtained after 10 (x-axis) and 15 (y-axis) minutes of 4sU labelling in wildtype (B) and DICER null (C) cells. (D) Volcano plot showing the adjusted p-value (y-axis) as a function of the fold-change in degradation rate estimates, based on the 10 minutes pulse, between KO and WT cells (x-axis) for protein-coding genes (red), cytosolic (blue) and nuclear (grey) lncRNAs. Each point represents a transcript and horizontal dashed line the significance cut-off. (E) Cumulative distribution plot of the density of AGO2 clusters in the 3’unstralated regions of AGO2 bound mRNAs (AGO2 cluster>0) whose degradation rates were either significantly (n=711, red) or not significantly changed (n=1127, black) between KO and WT cells, based on the 10 minutes pulse estimates. (F) Distribution of the fold-change after 8 days of tamoxifen treatment in degradation rate (estimated based on the 10 minutes pulse) of mRNAs (n=29900, red), cytosolic (n=474, blue) and nuclear (n=2348, grey) lncRNAs, in KO relative to WT cells. Statistics: *-p<0.05, **-p<0.01 and ***-p<0.001.

Next, we identified genes whose degradation rate is significantly different between miRNA-depleted and control mESCs (10 and 15 minute pulse, Figure 2D and Supplementary Figure S2E, respectively) and found that as expected, mRNAs are significantly more often stabilized in miRNA depleted mESCs relative to control. Finally, and consistent with a role of miRNA in controlling the observed differences in degradation rates, transcripts whose decay rates are significantly decreased, following miRNA depletion, have a significantly higher density of miRISC clusters (10 and 15 minute pulse, Figure 2E and Supplementary Figure S2F, respectively).

In contrast with mRNAs and in line with the observed changes in steady state abundances, we found that the degradation rates of cytosolic lncRNAs are minimally impacted by miRNA depletion, with only a few displaying significant differences in degradation rate (10 and 15 minutes pulse, Figure 2D and Supplementary Figure 2E). Specifically, most cytosolic lncRNAs behave similarly to nuclear lncRNAs (10 and 15 minutes pulse, Figure 2F and Supplementary Figure 2G, respectively). The decrease in degradation rate between wild-type and miRNA depleted cells is likely to be, at least in part a consequence of well described compensation mechanisms [37–39] to account for decreased synthesis rates between the two cell types (Supplementary Figure S2H). The analysis of steady-state abundance and degradation rates following loss of *Dicer* function indicate that, in contrast with coding transcripts, cytosolic lncRNAs are resilient to miRNA-mediated destabilization.

### Micropeptide encoding transcripts undergo miRNA dependent destabilization

Next, since our analysis of ribosomal profiling data indicated that a small fraction of cytosolic lncRNAs is ribosome-bound (Figure 1A), we investigated whether association with translating ribosomes would contribute to the impact of miRNAs on the degradation rates of some cytosolic lncRNAs. As expected, mRNAs are significantly more efficiently translated than lncRNAs but interestingly, the translation efficiency of cytosolic lncRNAs, as a class, is significantly higher than that of nuclear lncRNAs indicating that some might encode micropeptides (Figure 3A). The short open reading frames of micropeptide encoding transcripts are often missed by coding potential calculators leading to the misclassification of these transcripts as lncRNAs [40]. To distinguish *bonafide* lncRNAs from micropeptide encoding transcripts we used phyloCSF [41] and identified 59 cytosolic transcripts containing mammalian conserved short open reading frames (median longest predicted ORF length 216 nucleotides, Supplementary Table S1). These transcripts are almost 3 times more likely to be bound by ribosomes than are other cytosolic lncRNAs (Figure 3B) and their translation efficiency is significantly higher than that of cytosolic lncRNAs (*p*<6X10^-5^, two-tailed Mann-Whitney *U* test, Figure 3C) and more similar to that of mRNAs (*p*<1X10^-4^, two-tailed Mann-Whitney *U* test, Figure 3C) consistent with some of these transcripts encoding micropeptides. We separated micropeptides from *bona fide* cytosolic lncRNAs and found that fold change in degradation rate of micropeptides in miRNA depleted cells relative to control, is similar to what is obtained for mRNAs and significantly different from what is observed for *bonafide* lncRNAs (Figure 3D) indicating further the requirement of translation for miRNA-dependent transcript destabilization.

**Figure 3.**
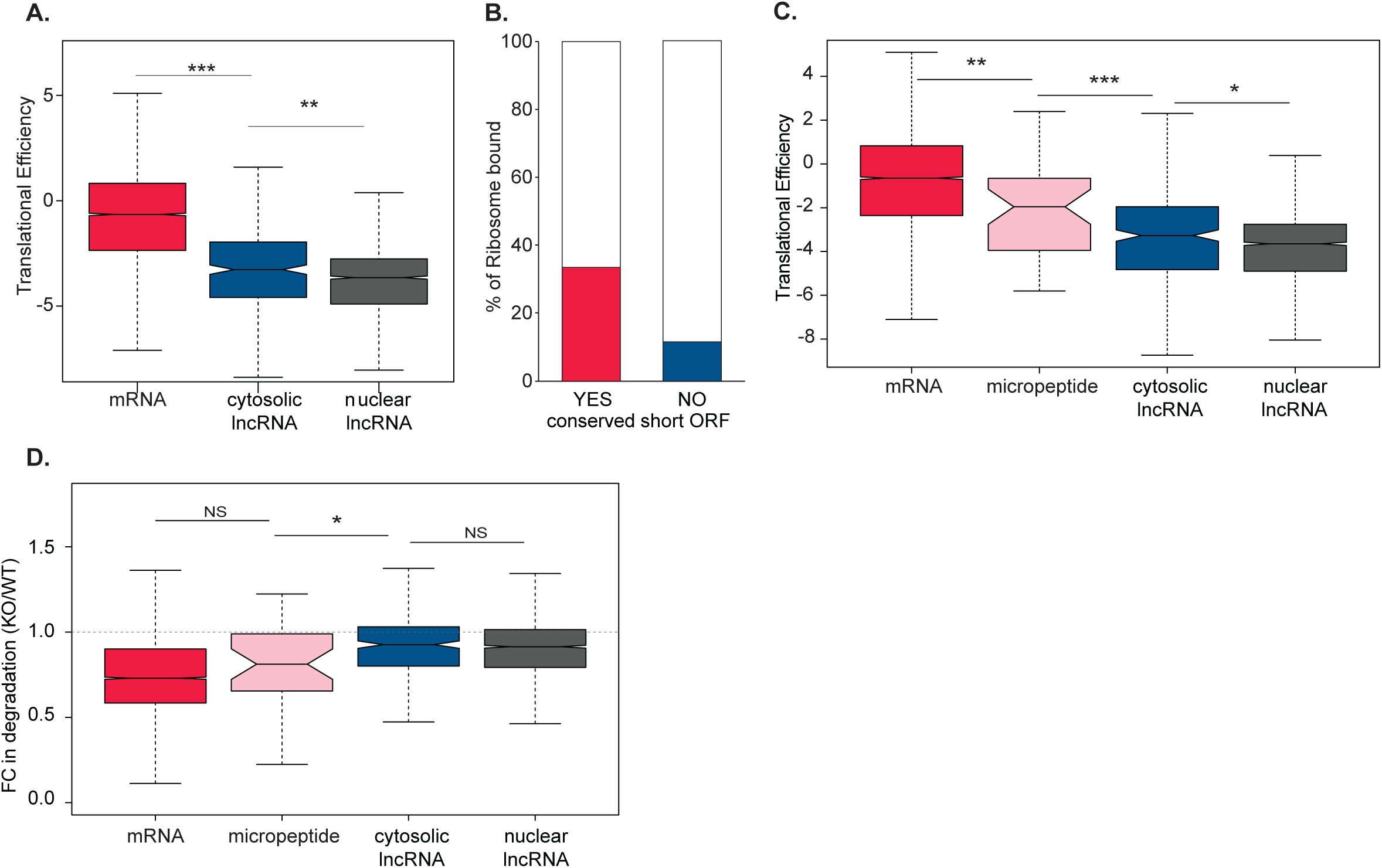
Micropeptide encoding transcript expression is posttranscriptionally regulated by miRNAs. (A) Distribution of the translational efficiency, in mESCs, of mRNAs (n=7156, red), cytosolic (n=341, blue) and nuclear (n=1915, grey) lncRNAs. Fraction of cytosolic lncRNAs with experimental evidence for ribosomal binding with (red) or without (blue) an overlapping conserved short open reading frame. (C) Distribution of the translational efficiency, in mESCs, of mRNAs (n=7156, red), micropeptide encoding transcripts (n=43, pink) and *bona fide* cytosolic (n=298, blue) and nuclear (n=1857, grey) lncRNAs. (D) Distribution of the fold-change after 8 days of 4-OHT treatment (KO) in degradation rates for mRNAs (n=13296, red), micropeptide encoding transcripts (n=43, pink) and *bona fide* cytosolic (n=759, blue) and nuclear (n=4299, grey) lncRNAs, relative to WT cells. Statistics: *-p<0.05, **-p<0.01 and ***-p<0.001.

### miRNA impact coding but not noncoding transcript stability

Our transcriptome wide analysis indicates that translation is required for miRNA dependent target destabilization. To test this hypothesis, we selected one cytosolic lncRNA (TCONS_00034281, Supplementary Figure S3A, hereafter lncRNA-c1) that is relatively highly expressed in mESCs (Supplementary Figure S3B). Quantitative PCR analysis supported that as indicated by the transcriptome wide profiling (Supplementary Figure S3C), the steady state abundance of *lncRNA-c1* does not increase upon miRNA depletion, as would be expected for *bonafide* miRNA target such as *Lats2* or *Cdkn1A* [35] (Supplementary Figure S3D). Furthermore, and in contrast with *Lats2* or *Cdkn1A*, *lncRNA-c1*’s stability is also not significantly affected in cells lacking DICER function (Supplementary Figure S3E). This is despite, lncRNA-c1 cytosolic localization (Supplementary Figure S3F) and binding by AGO2 that was suggested by AGO2-CLIP data and confirmed by AGO2-RIP (Supplementary Figure S3G-H).

We reasoned that if translation is required for miRNA-dependent transcript destabilization, forcing association of a lncRNA candidate to translating ribosomes, by fusing it downstream of a functional open-reading frame, should result in miRNA-dependent degradation of the fused transcript (Figure 4A). We cloned *lncRNA-c1* downstream of the Enhanced Green Fluorescent Protein stop codon (hereafter *GFP-lncRNA-c1*) and transfected this construct into wild-type and miRNA depleted mESCs (8 days after induction of *Dicer* loss of function). As controls, we transfected *GFP* and *lncRNA-c1* expressing constructs. As expected, the expression of *lncRNA-c1* and *GFP* is more similar between wild-type and miRNA-depleted cells than is the expression of *GFP-lncRNA-c1,* whose levels significantly increase in miRNA depleted cells (paired two-tailed t-test p-value < 0.02, Figure 4B), consistent with its miRNA dependent destabilization in wild-type cells. If association with the translation machinery is sufficient to induce miRNA-dependent decay of a miRISC-bound noncoding transcript, one would expect introduction of a missense mutation in a protein-coding miRNA target to decrease its miRNA-induced decay. Indeed, introduction of a missense mutation disrupting the *Cdkn1a* start codon (Supplementary Figure S4A-C) significantly decreases mutant *Cdkn1aΔATG* levels in miRNA depleted mESCs (paired one-tailed t-test p-value < 0.05, Figure 4C).

**Figure 4.**
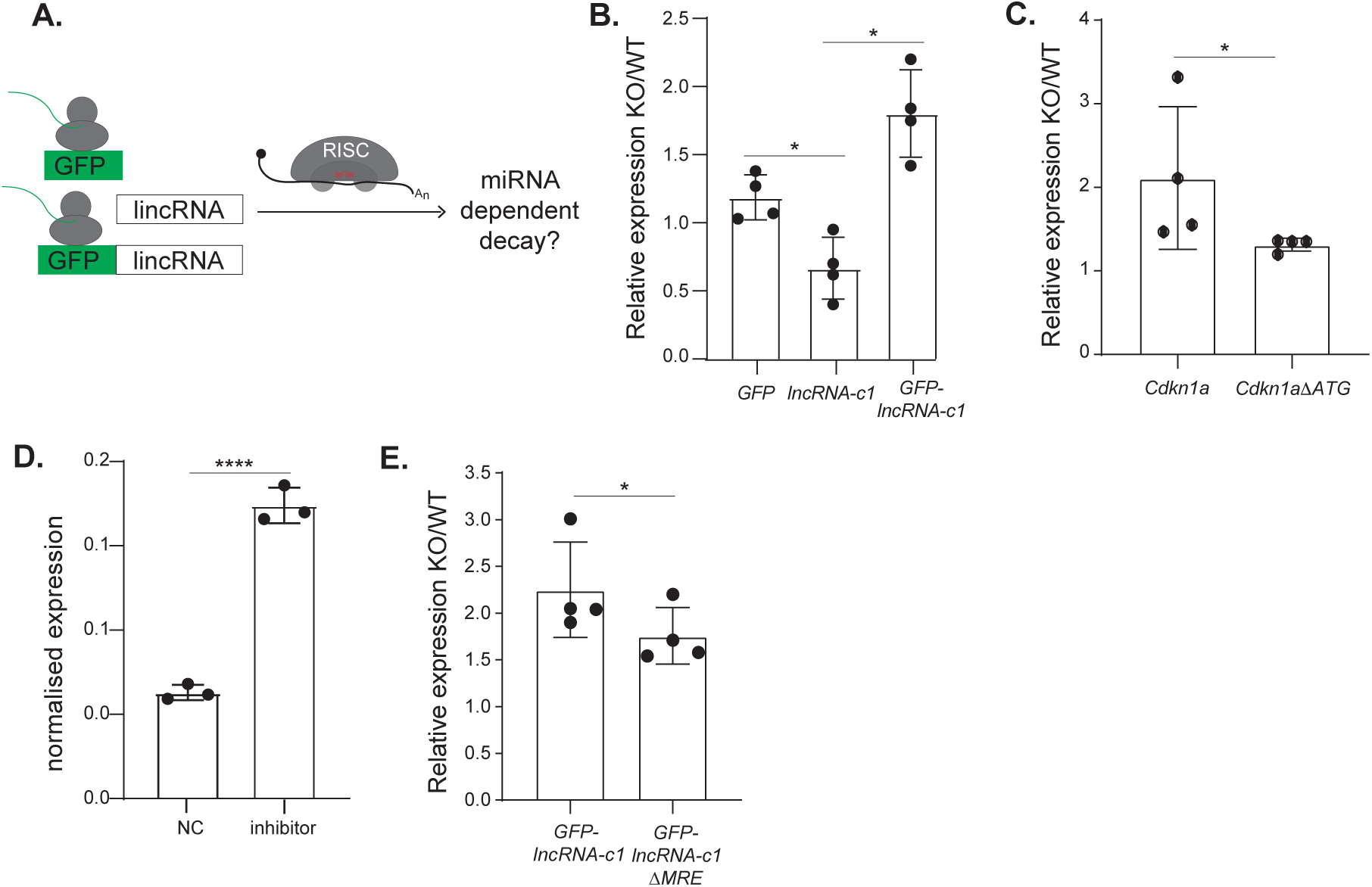
Association of lncRNA-c1 with translating ribosomes results in its miRNA-dependent decay. (A) Schematic of the construct tested in WT and miRNA depleted mESCs (B) Expression of *GFP*, *lncRNA-c1* and *GFP-lncRNA-c1* (x-axis) in miRNA depleted cells (KO) relative to WT mESC (y-axis) 24h hours post-transfection. Expression of *Cdkn1a and Cdkn1aΔATG* (x-axis) in miRNA depleted cells (KO) relative to WT mESC (y-axis) 24h hours post-transfection. (D) *GFP-lncRNA-c1* expression 24 hours following transfection of mESCs with miRNA294-3p inhibitors or small RNA negative control. (E) Expression of *GFP-lncRNA-c1* and *GFP-lncRNA-c1ΔMRE* (x-axis) in miRNA-depleted cells (KO) relative to WT mESC (y-axis). Transcript expression was first normalized by the amount of *Act-β* and *PolII* and next by the total amount of transfected vectors per cell estimated based on the levels of relative *Neomycin* expression. Each point corresponds to the results of one independent biological replicate. Statistics: *-p<0.05, **-p<0.01 and ***-p<0.001 two-tailed paired t-test.

Given that all constructs are under the control of the same promoter (T7), this increase is likely a consequence of increased stability, as confirmed by qPCR analysis following 8h of transcription inhibition through actinomycin-D treatment (paired two-tailed t-test p-value < 0.05, Supplementary Figure S4D).

*LncRNA-c1* is a predicted target of the miR-290/295 family (Supplementary Figure S4E). To validate that these miRNAs are indeed contributing to miRNA-dependent repression of *GFP-lncRNA-c1*, we co-transfected mESCs with *GFP-lncRNA-c1* expressing vector and miR-294-inhibitors. We note a significantly higher expression of *GFP-lncRNA-c1* in the inhibitor transfected cells compared to cells transfected with negative control (unpaired two t-test p-value < 0.001, Figure 4D). We used site-directed mutagenesis to mutate three miRNA recognition elements (MREs) for highly expressed miRNAs within *GFP-lncRNA-c1* (hereafter *GFP-lncRNA-c1ΔMRE*). As expected, reintroduction of miRNA mimics in DICER depleted mESC impacts the levels of wild-type *GFP-lncRNA-c1* more than it does levels of *GFP-lncRNA-c1ΔMRE* (paired one-tailed t-test p-value<0.05, Supplementary Figure S4F). The levels of *GFP-lncRNA-c1* in wild-type mESC is also significantly lower than the level of *GFP-lncRNA-c1ΔMRE* (paired t-test p-value<0.05, Supplementary Figure S4G). These results are consistent with these MREs’ contribution to wild-type *GFP-lncRNA-c1* miRNA-dependent repression. Therefore, and as expected, the relative increase of *GFP-lncRNA-c1* levels in miRNA depleted mESCs relative to wild-type mESC is significantly higher than the increase in levels of *GFP-lncRNA-c1ΔMRE* (paired two tailed t-test p-value < 0.05, Figure 4E). The presence of MRE for other mESC expressed miRNA (Supplementary Table S2) is likely to explain why mutation of miR290/295 MRE alone is not sufficient to entirely block miRNA-dependent *GFP-lncRNA-c1* destabilization.

We conclude that association with translating ribosomes is required for miRNA-dependent transcript destabilization and that noncoding transcripts are bound but not post-transcriptionally regulated by miRNAs.

## CONCLUSION

Posttranscriptional regulation by miRNAs leads to translational inhibition or transcript destabilization [6]. Whereas the general consensus is that most miRNA-induced changes can be explained by transcript destabilization [9, 10], increasing evidence suggests that miRNA-dependent mRNA decay occurs co-translationally [13–22], raising questions about the ability of miRNAs to posttranscriptionally regulate the levels of noncoding transcripts.

Supporting different outcomes upon miRISC binding to coding and noncoding transcripts, is recent evidence that these two classes of transcripts have distinct interaction dynamics with processing bodies (PB) [42], the subcellular compartment where miRNA-dependent destabilization is thought to occur [43]. Specifically, and in contrast with miRNA-bound mRNAs, which localise to the core of PB, miRNA-bound lncRNAs interact transiently and tend to locate to the PB periphery, a pattern that might reflect missing interactions with other molecular factors involved in miRNA-dependent regulation [42]. One such factor could be DDX6, a PB localised dead box helicase that links miRNA-dependent translation inhibition and decay [44–46]. In mESCs, loss of DDX6 function phenocopies loss of miRNA biogenesis [10, 47], suggesting that molecular factors that couple translation with RNA decay, like DDX6, are required for miRNA-dependent transcript destabilization.

These observations are surprising in light of previous analysis demonstrating efficient miRNA dependent decay in the absence of translation initiation or elongation [23–25]. One potential confounder of previous studies is that they rely on the use of exogenous reporters, which may not faithfully recapitulate what happens to endogenously expressed miRNA targets. Cytosolic *bonafide* lncRNAs, that have been previously shown to interact with miRISC [27] but not with the translation machinery [26], provide a unique opportunity to investigate the requirement of translation to endogenous miRNA-directed target decay.

Our transcriptome wide analysis following miRNA loss revealed, that in contrast with mRNA, cytosolic lncRNA’s steady state abundance significantly decreases in miRNA depleted cells, suggesting this class of transcripts is not efficiently posttranscriptionally regulated by miRNAs. To assess the direct impact of miRNA regulation on cytosolic lncRNAs, we investigated, using RNA metabolic labelling, differences in the degradation rates of these transcripts in wild-type and miRNA-depleted cells. This analysis revealed that cytosolic lncRNAs degradation rates decrease less than the degradation rates of mRNAs and to a similar extent as the degradation rates of nuclear lncRNAs, that are not expected to be regulated by miRNAs. The decrease of lncRNA degradation rates in miRNA depleted cells is likely the result of coupling between RNA synthesis and decay which has been proposed as a mechanism to ensure gene expression homeostasis [37–39]. While the decrease in degradation rates is a general phenomenon in miRNA-depleted mESCs (Supplementary Figure 2H), the increased stabilization of coding transcripts in near-absence of miRNA is likely to obscure such effects for mRNAs.

Finally, we show that the stabilities of putative micropetides and mRNAs are similarly impacted by miRNAs, further supporting the requirement of translation for miRNA dependent regulation of endogenously expressed transcripts.

To validate this hypothesis, we selected one cytosolic lncRNA, bound by AGO2 and with functional binding sites for miR-290/5 family, and forced its association to translating ribosomes by cloning it downstream of a functional protein-coding open reading frame. Consistent with the requirement of translation for miRNA-dependent transcript destabilization, forcing association to the ribosomes results in miRNA-dependent posttranscriptional regulation of previously unaffected transcripts. These results are unlikely a consequence of pleotropic effects of loss of miRNA function as mutation of the functional MREs within candidate lncRNA sequence reduces the impact of miRNAs on candidate expression. We conclude that miRNA-dependent regulation of endogenously expressed transcripts requires translation.

The requirement of translation for miRNA dependent regulation indicates that despite extensive evidence for miRISC binding to cytosolic lncRNAs, the levels of these noncoding transcripts are not posttranscriptionally modulated by miRNA. Evidence that miRNA binding sites within lncRNAs evolved under constraint [28] suggests that miRNA-lncRNA interactions are biologically relevant. One possibility, is that such interactions reflect miRNA-dependent regulation by lncRNAs. A number of examples support these roles in the context of disease and development [48–50]. Previous analysis of the potential extent of such regulatory roles by miRNAs suggested this mechanism of lncRNA function is prevalent among cytosolic transcripts [28]. However, given the relatively low abundance of most lncRNAs, which rarely exceeds the expected threshold to exert significant and physiological relevant changes in miRNA targets [51–53] the biological relevance of miRNA dependent regulation by lncRNAs remains controversial. In light of the present results, that support a different outcome of miRNA interactions with mRNA or lncRNAs, further experiments are now needed to assess the generality of mRNA-based conclusions.

More generally the present results also imply that miRISC binding, per se, is not sufficient to determine the outcome of bound targets suggesting the requirement of further yet unidentified molecular partners.

In summary, the analysis of endogenously expressed and miRISC bound noncoding transcripts provides further evidence that translation is indispensable for miRNA-dependent regulation of endogenous transcripts, suggesting the requirement of further molecular partners and highlighting differences in posttranscriptional regulation of coding and noncoding RNAs.

## METHODS

### KEY RESOURCES TABLE

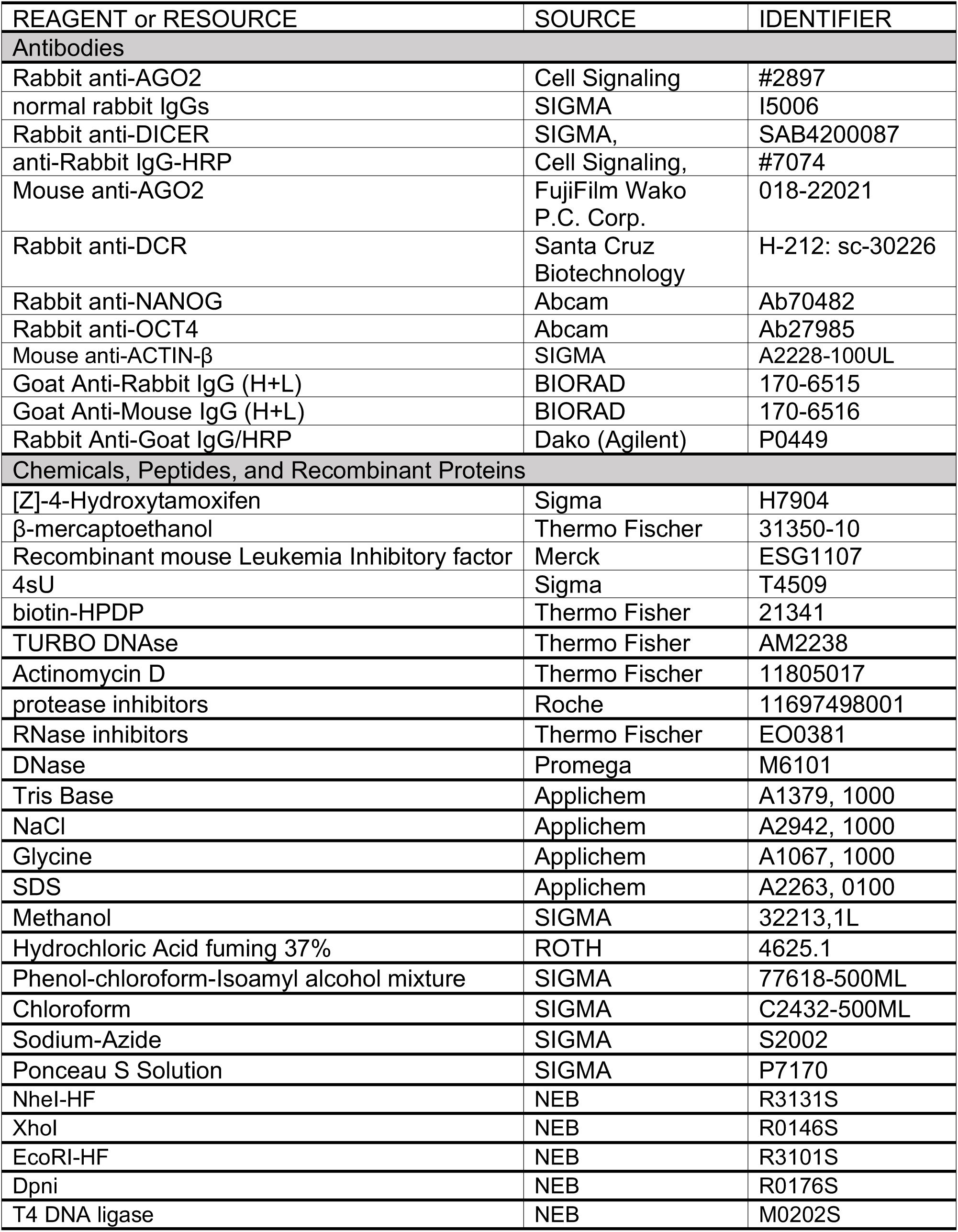
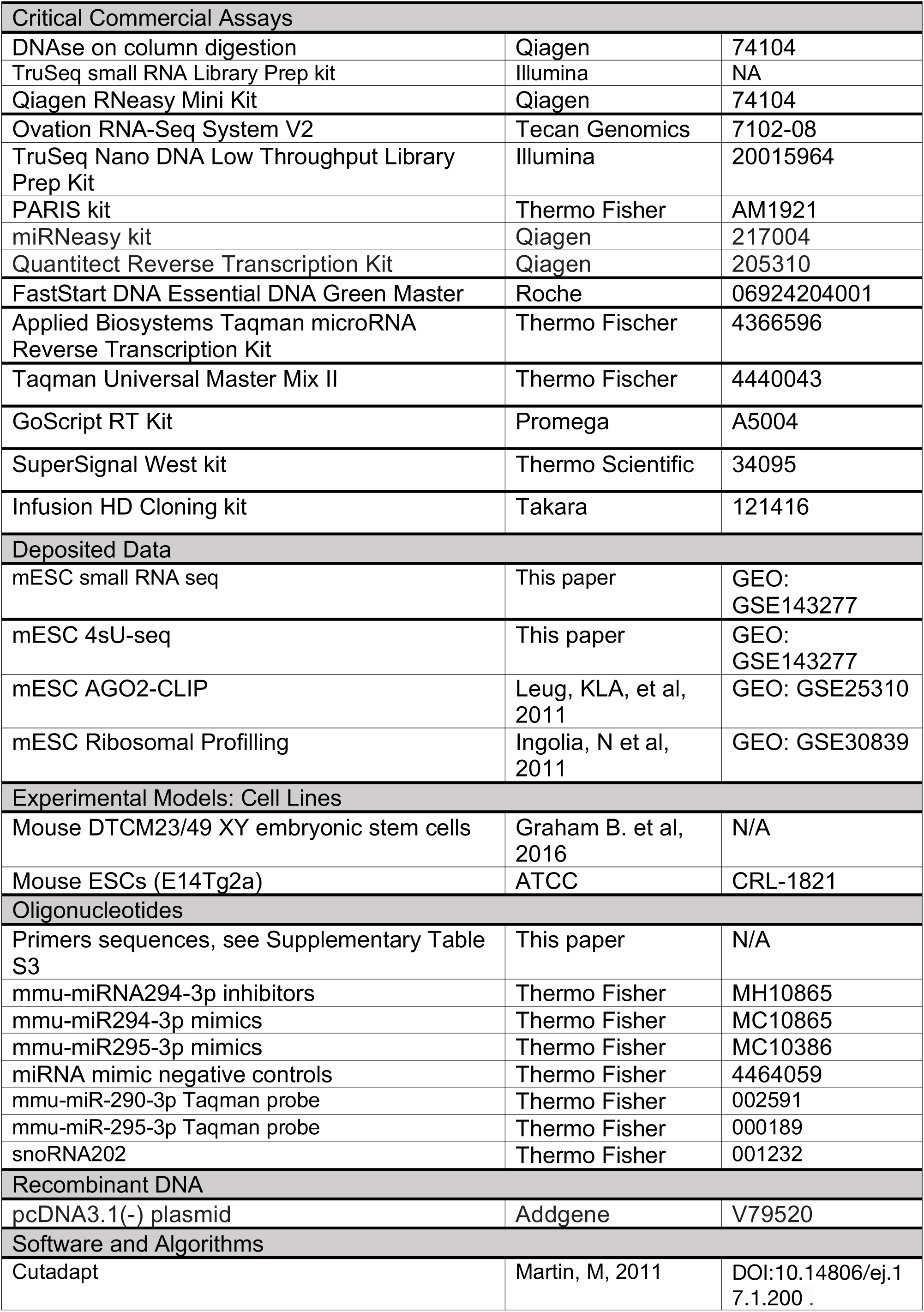
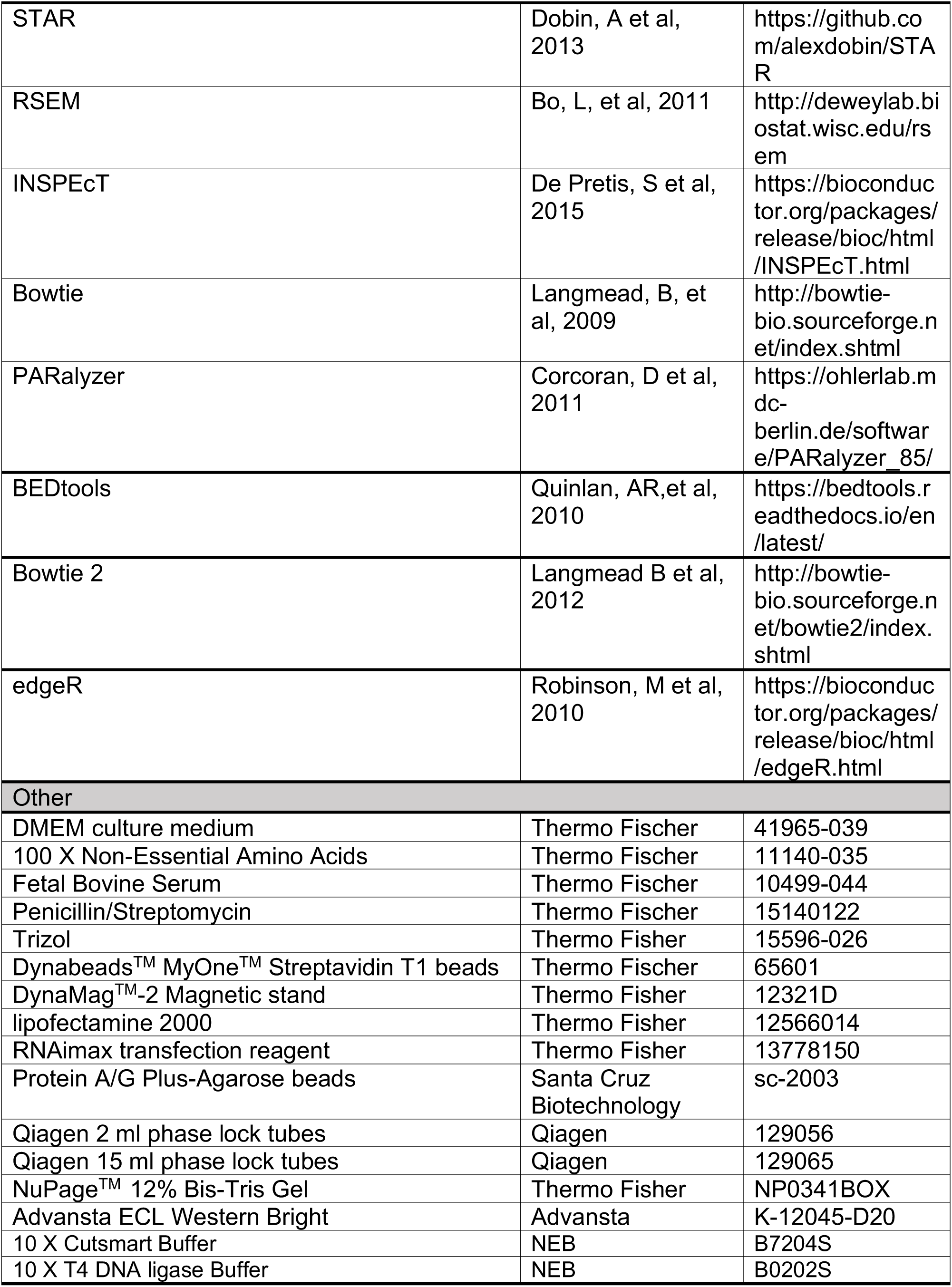

### Mouse embryonic stem cell culture

Feeder depleted mouse DTCM23/49 XY embryonic stem cells [28, 32, 54] were grown on 0.1% gelatin-coated tissue culture treated plates in a humidified incubator with 5% (v/v) CO_2_ at 37°C in 1X DMEM medium supplemented with 1x Non-Essential Amino Acids, 50 uM β-mercaptoethanol, 15% Fetal Bovine Serum, 1% Penicillin/Streptomycin, and 0.01% of Recombinant mouse Leukemia Inhibitory factor. Cultures were maintained by passaging cells every 48 hours (replating density ∼ 3.8*10^4^ cells/cm^2^). Unless stated otherwise, to induce loss of *Dicer* function, cells were cultured in mESC growth media supplemented with 800 nM tamoxifen previously resuspended in 100% ethanol ([Z]-4-Hydroxytamoxifen [4-OHT]) for 48h. Subsequently, cells were transferred to non-supplemented mESC growth medium and cultured for 6 additional days.

WT E14 mESC line (129/Ola background) was cultured in Dulbecco’s Modified Eagle Media (DMEM), containing 15% of fetal bovine serum, 100 U/mL LIF, 0.1 mM 2-ß-mercaptoethanol and 1% Penicillin/Streptomycin, on 0.2% gelatin-coated plates in absence of feeder cells. The culture medium was changed daily and cells were grown at 37°C in 8% CO_2_

### Small RNA extraction in Dicer depletion timecourse

Feeder depleted mouse DTCM23/49 XY embryonic stem cells were cultured in mESC growth media supplemented with 800 nM tamoxifen 4-OHT. Small RNA extraction and DNAse treatment following 0, 4, 8, 10 and 12 days of 4-OHT treatment was performed using the Qiagen miRNEasy Mini Kit and Qiagen RNAse free DNAse according to manufacturer instructions.

### Small RNA sequencing, mapping and quantification

Small RNA libraries were prepared from 500 ng of total RNA using Illumina TruSeq small RNA protocol and sequenced on Illumina HiSeq 2500.

Sequencing adapters were removed from fifty nucleotides long single-end reads using cutadapt (v1.8) and mapped to mouse genome (mm10) using bowtie2 (v2.2.4). Gene expression levels for all mouse miRNAs annotated in miRbase (v21) [55] were quantified using HT-seq (v0.6.1). The raw sequencing data and reads counts are available on the NCBI Gene Expression Omnibus (GEO) under accession number GSE143277.

### Western blot

Approximately 500,000 mESCs were harvested and washed twice with ice-cold PBS and stored, after PBS removal, at −80 °C until lysis. Cells were incubated in 50 µl of cold RIPA Buffer (150 mM NaCl, 1.0% NP-40, 0.5% sodium deoxycholate, 0.1% SDS, 50 mM Tris, pH 8.0) on a rotating wheel for 1 hour at 4°C. Protein concentration was determined using the Pierce BCA Protein Assay kit according to manufacturer’s instructions.

30 μl of protein was separated on NuPage^TM^ 12% Bis-Tris gel and transferred overnight at 4°C in transfer buffer (25 mM Tris-HCl pH7.6 192 mM glycine, 20% Methanol) on to nitrocellulose membranes. Transfer efficiency was assessed by staining the membrane with Ponceau S solution and staining solution was removed by washing the membrane 3 times with TBS-T (Tris-buffered saline, 0.1% Tween 20, 5 minutes, room temperature) After incubation with 5% skim milk in TBS-T for 4-6 hours at 4°C, the membranes were washed once in TBS-T and incubated with anti-DICER (1:3000), anti-NANOG (1:1000) or Anti-OCT4 (dilution 1:1000) antibodies in 5% skim milk in TBS-T overnight at 4°C on a see-saw shaker. After probing for protein of interest, membranes were stripped and probed for ACTIN-B as a loading control: Anti-ACTB (1:10000 dilution in 5% skim milk in TBS-T) Membranes were incubated with Secondary antibodies (DICER= 1:3000 Goat Anti-Rabbit IgG (H+L)-HRP Conjugate; for NANOG= 1:4000 Goat Anti-Rabbit IgG (H+L)-HRP Conjugate; for OCT4=1:2500 Rabbit Anti-Goat IgG/HRP, for ACTIN-B=1:2000 Goat Anti-Mouse IgG (H+L)) in 5% skim milk in TBS-for 1h at room-temperature. Immunoblots were developed using the WesternBright ECL premixed Peroxide and ECL solutions and detected using an imaging system (Vilber Fusion Chemiluminescence).

Following detection, secondary antibody coupled with the HRP was deactivated by washing the membrane two times for 20 minutes with 1% (w/v) Sodium-Azide in TBS-T and the membrane incubated two hours at 4 °C with 5% (w/v) skimmed milk in TBST containing primary antibody for the ACTIN-β loading control (1:4000). The membranes were washed three times for 15 minutes in fresh TBS-T and incubated for one hour at room temperature with the secondary antibody coupled with horseradish peroxidase in 5% skimmed milk in TBS-T (for ACTIN-β= 1:4000 Goat Anti-mouse IgG). Washing and protein detection were performed as previously described.

### Cell proliferation assay

16-24 hours prior to DNA staining, 33,000 cells/cm^2^ were plated on a 6-well gelatin-coated tissue culture plate. Edu (Click-iT Edu Alexa Fluor^TM^ 488 Flow Cytometry Assay Kit) was added to mESCs growth medium to a final concentration of 10 µM, and the cells incubated at 37 °C for 30 minutes. Cells were trypsinized, counted and, for each tested sample, 750,000 cells were washed once with 3 ml of 1% BSA in PBS, resuspended in 100 µl of Click-iT fixative buffer and incubated for 15 minutes at room temperature in the dark. Cells were washed with 3 ml of 1%BSA in PBS, centrifuged and the supernatant removed. The pellet was resuspended in 100 µl of 1X Click-iT saponin-based permeabilization and wash reagent, and the cells incubated for 15 minutes at room temperature in the dark. 500 µl of freshly prepared Click-iT reaction cocktail containing Alexa Fluor 488 Fluorescent dye Azide was added to the permeabilized cells in 1X Click-iT saponin-based permeabilization and wash reagent and the mix incubated at room temperature in the dark for 30 minutes. Cells were washed once with 3 ml of 1X Click-iT saponin-based permeabilization and wash reagent and following supernatant removal resuspended in 500 µl of Click-iT saponin-based permeabilization and wash reagent. Cells were analyzed by flow cytometry on a Beckman Coulter Gallios Flow Cytometer according to manufacturer’s instructions, using a 488 nm excitation wavelength and a green emission filter (530/30 nm).

### 4sU metabolic labelling

Five million DTCM23/49 XY mESCs (WT and miRNA depleted) were seeded and allowed to grow to 70-80% confluency (approximately 1 day). 4sU was added to the growth medium (final concentration of 200 µM) and cells were incubated at 37 °C for 10 or 15 minutes. RNA was extracted using Trizol, according to manufacturer instructions and DNAse treated using RNeasy on column digestion according to manufacturer’s instructions. 100 µg of RNA was incubated for 2 h at room temperature with rotation in 1/10 volume of 10X biotinylation buffer (Tris-HCl pH 7.4, 10 mM EDTA) and 2/10 volume of biotin-HPDP (1mg/ml in Dimethylformamide). Following biotinylation, total RNA was purified through phenol:chloroform:isoamyl alcohol extraction and precipitated with equal volume of Isopropanol and 1/10 volume of 5M NaCl. RNA was washed once with 75% Ethanol and resuspended in in DEPC-treated H_2_O. Equal volume of biotinylated RNA and pre-washed Dynabeads^TM^ MyOne^TM^ Streptavidin T1 beads were mixed and incubated at room temperature for 15 minutes under rotation. The beads were then separated using a DynaMag^TM^-2 Magnetic stand. The supernatant (that contains unlabeled preexisting RNA) was placed at 4°C until precipitation. Beads were washed and biotinylated RNA dissociated from streptavidin coated beads by treatment with 100 mM 1,4-Dithiothreitol for 1 minute, followed by 5 minutes in RTL buffer. Beads were separated from the solution using DynaMag^TM^-2 Magnetic stand and the RNA recovered from the supernatant extracted using Qiagen RNeasy Mini Kit according to the manufacturer’s instructions. Preexisting RNA was precipitated with equal volume of Isopropanol and centrifuged for 45 minutes at 15 000 g at 4°C. Preexisting RNA pellet was washed with 75% Ethanol and resuspended in DEPC-treated H_2_O. Metabolic labelling experiments were repeated once for the 2 labelling durations (2 biological replicates).

### RNA sequencing, mapping, and quantification of metabolic rates

Total RNA libraries were prepared from 10 ng of DNase-treated preexisting and newly transcribed RNA using Ovation^®^ RNA-Seq and sequenced on an Illumina HiSeq 2500 (average of fifty million reads per library).

Hundred nucleotides long single-end reads were first mapped to *Mus musculus* ribosomal RNA (rRNA, ENSEMBL v91,[62]) with STAR v2.5.0 [56]. Reads that do not map to ribosomal RNA were then aligned to intronic and exonic sequences of *Mus musculus* transcripts database (ENSEMBL v91) using STAR and quantified using RSEM [57]. Principal Component Analysis (PCA) of read counts was performed to demonstrate separation between newly-transcribed (labeled) and total RNA (Figure S1D). Rates were inferred, independently at each labeling point using the INSPEcT ([35] Bioconductor package v1.8.0). Specifically, the absolute values of synthesis, processing and degradation rates in each condition were estimated using the ‘newINSPEcT’ function with the option pre-existing=TRUE, while the statistical significance of the variation of the rates between conditions was obtained using the method ‘compareSteady’ [see INSPEcT vignette at http://bioconductor.org/packages/INSPEcT/]. The raw sequencing data is available on the NCBI Gene Expression Omnibus (GEO) under accession number GSE143277.

### Identification of AGO2 bound regions in mESCs

Cutadapt [58] was used to remove sequence adapters from publicly available AGO2-CLIP sequencing reads from wild-type and *Dicer* mutant mESCs [30]. Trimmed reads were mapped to the mouse genome (mm10) using bowtie [59] (bowtie -v 2 -m 10 -- best –strata) as previously described [60]. Mapped reads from the same cell type were merged AGO2 bound clusters identified using PARAlyzer v1.5 (Bandwidth=3; minimum read count per group=5; minimum read count per cluster=1; minimum read count for KDE=5; minimum cluster size=1; minimum conversion count per cluster=1; minimum read count for cluster inclusion=1) [60]. Clusters present in wild-type and DICER null cells were excluded using BEDtools [61].

### Translational efficiency

Ribosome profiling (RP) and total RNA raw reads were downloaded from SRA database (SRX084815 and SRX084812, respectively [29]). Reads were trimmed based on quality and sequence adapters removed with Cutadapt (v. 1.8,[58]). Only reads with the expected read length (16 to 35 nt for the ribosome footprint and 35 to 60 nt for total RNA) were kept for further analysis. Reads mapping *Mus musculus* ribosomal RNA (rRNA) and transfer RNA (tRNA) databases (ENSEMBL v91,[62]) using to bowtie2 (v. 2.3.4.1, parameters: -L 15 -k 20,[63]) were excluded. The remaining reads (SRX084815: 12 228 002 reads; SRX084812: 12 361 681 reads) were aligned against *Mus musculus* transcripts database (ENSEMBL v91) using bowtie2 (v. 2.3.4.1, -L 15 -k 20). Multi-mapping reads (mapping to 2 or more transcripts from different gene loci) were filtered out and the remaining reads summarised at a gene level using an in-house script. Translational efficiency (TE) was calculated in R. Briefly, raw genes ribosome footprints and total RNA counts were normalized using the edgeR package to account for variable library depths (cpm function; [64]).Translational efficiency (TE) was calculated as the log2 ratio between normalized RP counts and normalized TR counts.

Conserved short open reading frames within lncRNA transcripts were identified by overlapping lncRNA loci with regions with positive phyloCSF scores, those that likely represent conserved coding regions, in any of the three possible reading frames on the same strand as the lncRNA transcript ([41]). LncRNA transcripts containing conserved short open reading frames are likely to encode micropeptides.

### Subcellular Fractionation

Subcellular fractionation of mESCs was carried out using the PARIS kit according to the manufacturer’s instructions. Following RNA extraction from cytosolic and nuclear fractions, genomic DNA was removed from samples using TURBO DNAse. DNAse treated RNA was extracted using phenol chloroform and RNA precipitated using equal volume of isopropanol and 1/10 volume of 5M NaCl. RNA pellet was washed with 75% Ethanol and resuspended in DEPC-treated H_2_O.

### RNA extraction and qPCR

Total cellular RNA was extracted with the RNeasy Mini kit according to the manufacturer’s instructions. To quantify levels of mature miRNAs, total RNA was extracted with the miRNeasy kit. Genomic DNA was removed by performing an on column DNAse treatment according to manufacturer’s instructions. Following RNA elution in DEPC-H_2_O, an additional DNAse treatment was performed using TURBO DNAse as described above. Following precipitation, RNA was reverse transcribed using the Quantitect Reverse Transcription Kit. Quantitative PCR reactions were prepared using the FastStart DNA Essential DNA Green Master and sequence-specific primers (Supplementary Table S3 and analyzed using a Roche Light Cycler®96. Unless otherwise stated *Actin-β* and *PolymeraseII* were used as internal controls.

For miRNA level quantification, RNA was reverse transcribed using the Applied Biosystems Taqman microRNA Reverse Transcription Kit and small RNA specific probes according to manufacturer’s protocol. Small RNA expression levels relative to *small nucleolar RNA 202* (sno-202) were subsequently quantified on a Roche Light Cycler®96 using the Taqman Universal Master Mix II, no UNG, according to manufacturer’s instructions.

### RNA stability

Transcription was inhibited by adding Actinomycin D resuspended in Dimethyl Sulfoxide at a final concentration of 10 µg/ml in supplemented mESC growth medium. Stability of transcripts was inferred by comparing relative gene expression levels (normalized to *Actin-β*) in cells incubated for 8 hours with Actinomycin-D and untreated cells.

### Candidate lncRNA and mRNA analysis

Enhanced Green Fluorescent Protein gene (see Supplementary Table S3) was amplified from the pBS-U6-CMV-EGFP plasmid [65] with primers complementary to EGFP and NheI restriction sites (see Supplementary Table S3) and inserted into NheI digested pcDNA3.1(-)(Addgene, V79520). Ligation was performed using T4 DNA ligase according to manufacturer instructions. Plasmid was transformed into DH5α subcloning efficiency bacterial cells and Sanger sequencing was used to confirm correct orientation of EGFP insertion into plasmid (*GFP)*.

*lncRNA-c1* was amplified from mESC cDNA using sequence specific primers with overhangs containing restriction sites for either XhoI or EcoRI (Supplementary Table S3) and cloned directionally into XhoI-EcoRI digested pcDNA3.1(-) plasmid to generate *lncRNA-c1* construct downstream of T7 promoter. Ligation was performed using T4 DNA ligase according to manufacturer instructions. *GFP-lncRNA-c1* construct was generated adopting same cloning strategy but inserting *lncRNA-c1* into *GFP* construct. Sanger sequencing was used to confirm correct sequence.

*Cdkn1a* was amplified from mESC cDNA using sequence specific primers with overhangs containing restriction sites for either XhoI or EcoRI (Supplementary Table S3) and cloned directionally into XhoI-EcoRI digested pcDNA3.1(-) plasmid downstream of T7 promoter. Forward primers *Cdkn1aΔATG* introduce a missense mutation that deletes the 1^st^ position of the *Cdkn1a* start codon (Supplementary Table S3). Ligation was performed as previously described and the correct sequence of constructs was confirmed by Sanger sequencing.

2 MRE on *GFP-lncRNA-c1* were mutated using, the Takara In-fusion HD cloning kit according to manufacturer’s instructions. The primers were designed using the manufacturer online design tool (https://www.takarabio.com/learning-centers/cloning/in-fusion-cloning-tools) and are available in Supplementary Table S3. The MREs were mutated sequentially using primer containing the mutation of interest and AmpHiFi PCR Master Mix. PCR products were gel purified, ligated using the In-fusion HD enzyme and transformed into Stellar competent bacterial cells according to manufacturer’ instruction. MRE mutation was confirmed through Sanger sequencing.

1 MRE on *GFP-lncRNA-c1* was mutated using the Phusion High-Fidelity Polymerase. Briefly, primers containing a scrambled sequence of the seed region within the MRE and wings complementary to the targeted sequence (Supplementary Table S3) were used to amplify from the *GFP-lncRNA-c1* containing plasmid. Following amplification PCR purification was performed and DpnI digestion was used to digest template plasmid. Blunt end ligation using T4 DNA ligase was performed to ligate amplified sequence containing mutated MRE according to manufacturer instructions. Ligated construct was subsequently transformed into DH5α bacterial cells and MRE mutation was confirmed through Sanger sequencing.

One day prior to transfection wild-type and miRNA depleted DTCM23/49 XY embryonic stem cells were plated in 10 cm dishes at a density of 35000 cells/cm^2^. Cells were transfected with 484×10^−15^ mol of candidate expressing vector using the lipofectamine 2000 transfection reagent. RNA was extracted 24 hours after transfection. Gene expression levels relative to *Actin-β* and *PolymeraseII* of transfected candidates were normalized to *Neomycin* expression to account for differences in transfection efficiency between different cell types and experiments. To distinguish between mRNA endogenous and exogenous expression, *Cdkn1a* levels were measured using Bovine Growth Hormone (BGH) polyadenylation signal specific primers (Supplementary Table S3) which bind upstream of the termination site of the exogenously expressed *Cdkn1a* constructs.

For miRNA mimic and inhibitor transfections, mmu-miRNA294-3p inhibitors (30 mM), mmu-miR294-3p, mmu-miR295-3p mimics (100 mM) and miRNA mimic negative controls were transfected 24 hours after plasmid transfection using the RNAimax transfection reagent according to manufacturer’s instructions. RNA was extracted 24 hours after small RNA transfection and reverse transcription was performed according to manufacturer’s instructions as described above.

### RNA immunoprecipitation

RNA immunoprecipitation was performed as previously described [66]. Briefly, 4.8×10^6^ E14 WT cells were seeded into 10cm dishes 16 hours prior to harvest. At the same time, 60µl of Protein A/G Plus-Agarose beads were incubated with 10µl of Rabbit anti-AGO2 or 2.5µg of normal rabbit IgGs. Protein content in cell lysate was split in half, adjusted to 1ml using IP Lysis buffer and supplemented with protease inhibitors and RNase inhibitors. 50µl of diluted cell lysates were collected for Input (5%). The remaining cell lysate was added to the A/B or IgG coupled beads and incubated overnight at 4° C, on a rotating wheel. After washing, 100 µl of RIP buffer + 1 µl of RNAse inhibitor was added to the beads and centrifuged. 20 μl and 80 μl of supernatant were collected for protein and RNA analysis. Immunoprecipitated and input RNA was extracted using TRIzol reagent, resuspended in DNAse reaction mix (16µl ddH_2_O, 2µl 10x RQ1 DNase buffer, 2µl RQ1 DNase) and reverse transcribed using the GoScript RT Kit and oligo d[T]_18_. RT-qPCR were performed as described above.

10 μl of RIP supernatant and input samples were separated on 8% SDS-PAGE gels and transferred to PVDF membranes. After incubation with 5% skim milk in 1xPBS/0.1% Tween-20, the membranes were washed and incubated with antibodies against AGO2 (ARGONAUTE 2 Rabbit mAb) and Dicer (Rabbit anti-Dicer) at 4° C. for 16h. Secondary antibodies (anti-Rabbit IgG-HRP) were incubated on membranes for 1h at RT at a dilution of 1:5000. Immunoblots were developed using the SuperSignal West kit and detected using an imaging system. Membrane stripping was performed by low pH method and AGO2 membrane was re-probed with antibodies against AGO2 (Argonaute 2 Mouse mAb). All membranes were stained using a coomassie blue staining solution to ensure equal loading.

### Data and Code Availability

RNA sequencing data was analyzed as described in Method Details; the data files are available in the Gene Expression Omnibus accession number GEO: GSE143277. Unprocessed Western blot images are available at Supplementary File 1 and 2.

## Acknowledgements

We would like to thank the Genomics Technology Facility at the University of Lausanne for help with RNA integrity analysis, library preparation and sequencing.

We would like to thank Maria Ferreira Da Silva for performing the Flow Cytometry analysis of the Proliferation Rate assay. The computations were performed at the Center for high-performance computing of the University of Lausanne.

This work was funded by the Swiss National Science Foundation (grant PP00P3_150667 to A.C.M and 31003A_173120 to C.C) and the NCCR in RNA & Disease (A.C.M. and C.C.).

## Authors Contributions

AB and ACM designed the study. AB, SdP, JYT, RD and ACM performed the in silico analysis. AB, BA, HW, CC and ACM performed and analyzed *in vitro* experiments analysis. MP, CC and ACM supervised the study. ACM wrote the manuscript. All coauthors read and approved the manuscript.

---

**Supplementary Table S1 - Gene IDs and locations of putative micropeptides.**

**Supplementary Table S2 - Location and identity of lncRNA-c1 predicted miRNA recognition sites for mESC expressed miRNAs.** Only miRNAs belonging to the 14 families that account for 75% of all miRNA counts, as estimated using nanostring in [28], were considered. MREs mutated in GFP-lncRNA-c1ΔMRE are highlighted in red.

**Supplementary Table S3 - Primer table.**

**Supplementary Figure S1.**
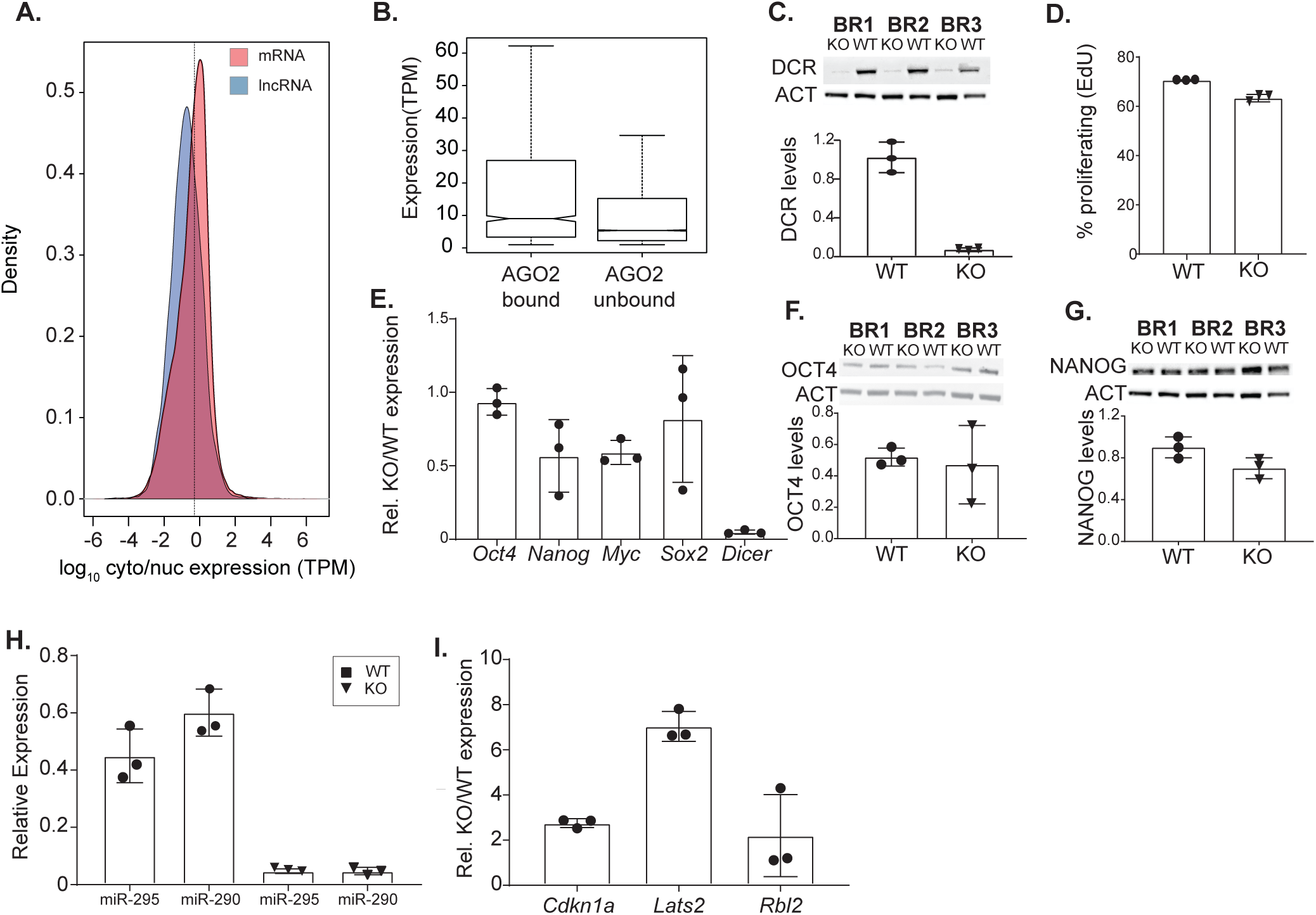
(A) Distribution of log 10 ratio between nuclear/cytosolic (transcripts per million, tpm) in mESCs for mRNAs (red) and lncRNAs (blue) (B) Distribution of the expression (tpm) of transcripts with and without experimental evidence for AGO2 binding in mESCs. (C) Immunoblot analysis of DICER (DCR) in protein extracts from DICER conditional mESCs 8 days after treatment with ethanol (WT) or tamoxifen (KO) for three independent biological replicates (BR1-3). ACTIN-β (ACT) was used as an internal control and to quantify the relative difference in DICER levels represented in bar plot. (D) Percentage of proliferating cells after 8 days of treatment with ethanol (WT) or 4-OHT (KO) for 3 independent treatments. (E) Fold-change in *Oct4* (two-tailed t-test p-value=0.48), *Nanog* (two-tailed t-test p-value=0.09), *Myc* (two-tailed t-test p-value=0.036), *Sox2* (two-tailed t-test p-value=0.38) and *Dicer* (x-axis) (two-tailed t-test p-value<0.0001) expression KO relative to WT cells measured for 3 independent biological replicates (y-axis). Western blot using Antibodies against mouse OCT4 (two-tailed t-test p-value=0.88) (F) and NANOG (two-tailed t-test p-value=0.70) (G) in protein extracts from DICER conditional mESCs 8 days after treatment with ethanol (WT) or 4-OHT (KO) for three independent biological replicates (BR1-3). ACTIN-β (ACT) was used as an internal control and to determine the relative difference in DICER levels represented in bar plot. (H) Expression of miR-295 and miR-290 relative to sno-202 (x-axis) in Dcr conditional mESCs 8 days after treatment with ethanol (WT) or tamoxifen (KO) (y-axis) for three independent biological replicates. (I) Fold-change in *Cdkn1a*, *Lats2* and *Rbl2* (x-axis) expression KO relative to WT cells measured for 3 independent biological replicates (y-axis). Uncropped blots used to assemble panels C, F and G are provided in Supplementary Files 1-2.

**Supplementary Figure S2.**
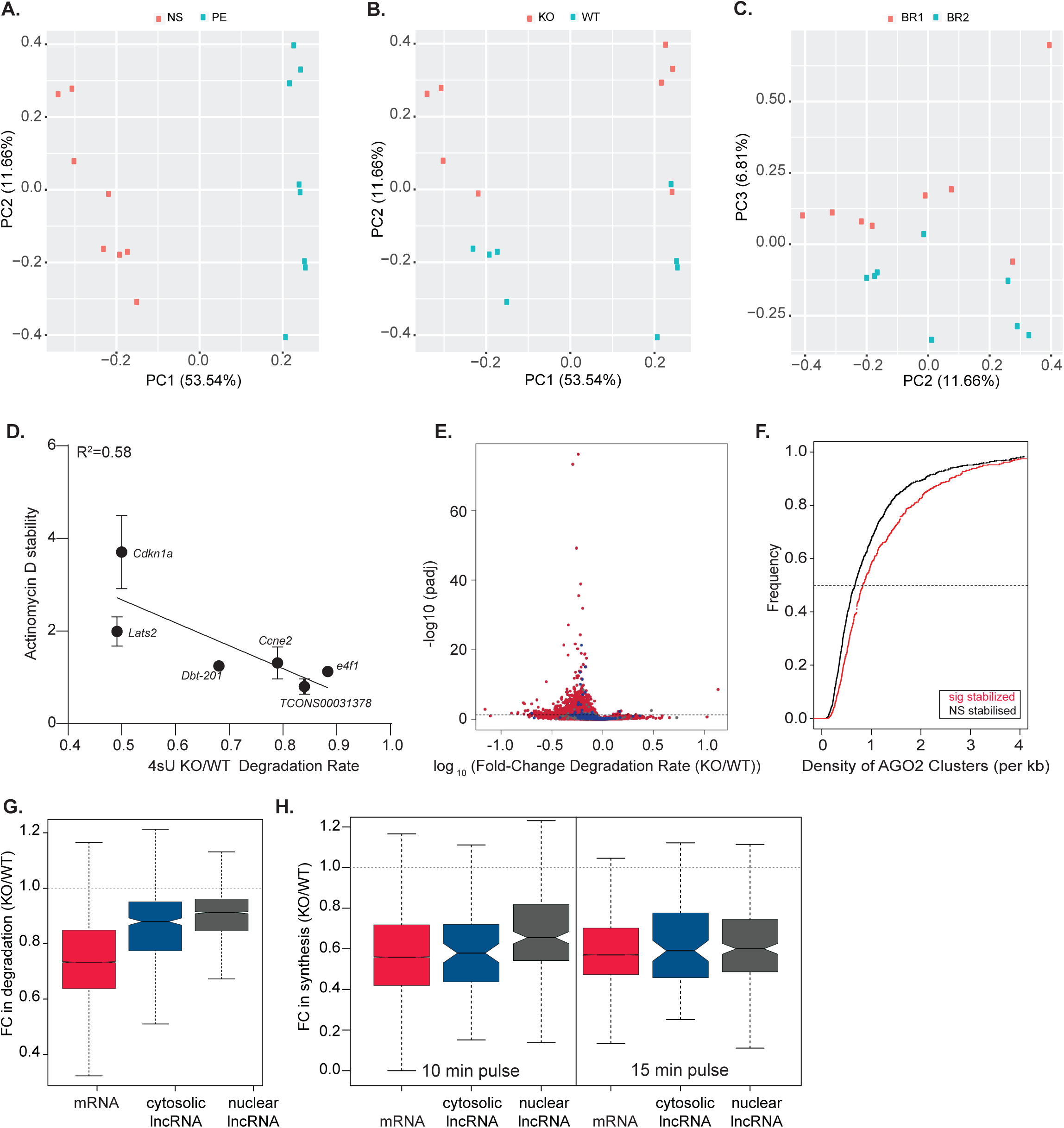
Principal component analysis of gene expression. The first 2 axis (PCA1 and PCA2) separate samples into (A) RNA fraction, nascent RNA (NS, red) and preexisting RNA (PE blue) and (B) cell type, DICER knockout (KO, red) and wild-type (WT, blue). (C) PCA2 and PCA3 separate biological replicates (BR1 red and BR2 blue). (D) Fold-change in 4sU degradation rate between KO and WT cells (X axis) is inversely correlated with the fold-change in relative expression between KO and WT after 8 hours of treatment with Actinomycin-D (y-axis). Points represent the mean and standard deviation based on 3 independent biological replicates. (E) Volcano plot showing the adjusted p-value (y-axis) as a function of the fold-change in degradation rate, estimates based on the 15 minutes pulse, between KO and WT cells (x-axis) for protein-coding genes (red), cytosolic (blue) and nuclear (grey) lncRNAs. Each point represents a transcript and horizontal dashed line the significance cut-off. (F) Cumulative distribution plot of the density of AGO2 clusters in the 3’unstralated regions of mRNAs bound (AGO2 cluster>0) whose degradation rates were either significantly (red) or not significantly changed (black) between KO and WT cells, based on the 15 minutes pulse estimates. (G) Distribution of the fold-change after 8 days of tamoxifen treatment in degradation rate (estimated based on the 15 minutes pulse) of mRNAs (red), cytosolic (blue) and nuclear (grey) lncRNAs, in KO relative to WT cells. (H) Distribution of the fold-change after 8 days of tamoxifen treatment in synthesis rate of mRNAs (red), cytosolic (blue) and nuclear (grey) lncRNAs, in KO relative to WT cells. Results for the 10- and 15-minutes pulse are presented separately.

**Supplementary Figure S3.**
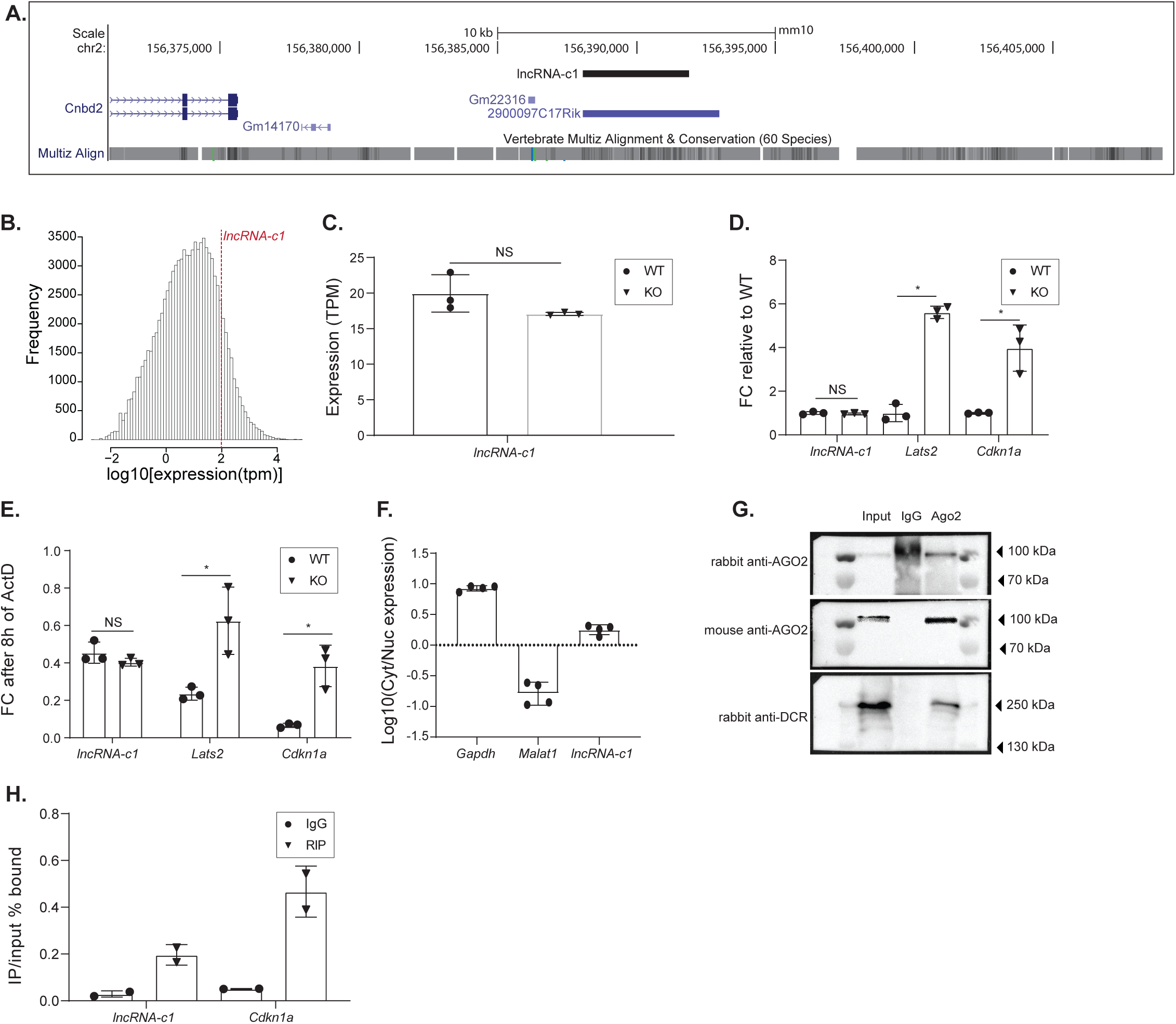
(A) Genome browser view of the region encompassing *lncRNA-c1* (black, chr2: 156388130-156391779). Gencode annotated genes are annotated in blue. (B) Distribution of gene expression (log10tpm, x-axis) for all mESC expressed transcripts. Red dotted horizontal line indicates the expression of lncRNA-c1. (C) Expression of *lncRNA-c1* (TPM), measured by RNA sequencing, 8 days after induction of DICER loss of function in wild-type (WT, circles) and 4-OHT treated (KO, triangles) mESCs. Each point represents the expression measured in one of the biological replicates. (D) Fold-change in *lncRNA-c1*, *Lats2* and *Cdkn1a* expression relative to WT cells, measured by qPCR in WT (circles) and 4-OHT treated (KO, triangles) mESCs. Transcript expression was normalized by the amount of *Act-β* and *PolII*. (E) Fold-change in stability, measured as the relative amount of transcript detected after 8 hours of transcription block using Actinomycin-D, for *lncRNA-c1*, *Lats2* and *Cdkn1a* expression relative to WT cells. Expression was measured by qPCR after in WT (circles) and 4-OHT treated (KO, triangles) mESCs for tree independent experiments. (F) Log10 of the fold change in expression in the nuclear and cytosolic fraction. Measured by qPCR, for *lncRNA-c1* and a nuclear (*Malat1*) and cytosolic (*Gapdh*) control. (G) Representative western blot analysis of protein extracts from input, AGO2-RIP and IgG control. AGO2 was probed with rabbit AGO2 antibody (top panel). After membrane stripping and re-probing with mouse anti-AGO2 (middle panel) unspecific band in IgG was cleared. Probing with rabbit antibody confirmed the presence of DICER specifically in the input and AGO2-RIP samples (lower panel). (H) qPCR quantification of *lncRNA-c1* and *Cdkn1a* (x-axis) bound in AGO2-IP (triangles) relative to input and unspecific IgG (circles) antibody relative to input (y-axis).

**Supplementary Figure S4.**
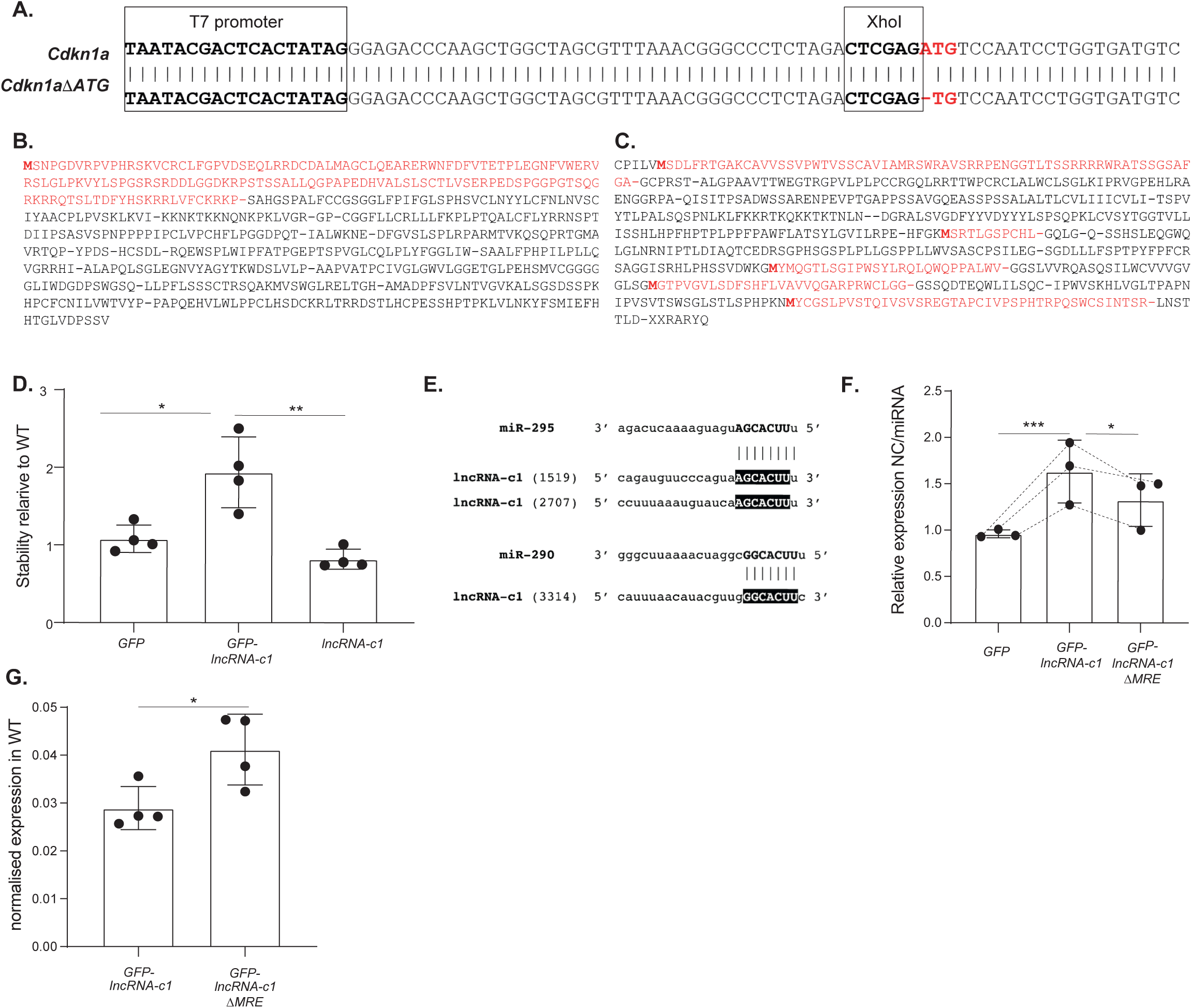
(A) Pairwise alignment of the constructs sequencing results *for Cdkn1a* (top) and *Cdkn1aΔATG* (bottom). *Cdkn1a* start codon is highlighted in red. Predicted peptides encode by (B) *Cdkn1a* and (C) *Cdkn1aΔATG* (same frame). (D)Fold-change in stability, measured as the relative amount of transcript, normalized to *Act-β,* detected after 8 hours of transcription block using Actinomycin-D, for *GFP*, *GFP-lncRNA-c1* and *lncRNA-c1* (x-axis) in miRNA depleted cells relative to WT cells (y-axis). (D) Pairwise alignment between miR-295 (top) and miR-290 (bottom) and respective predicted miRNA response elements (MRE) within *lncRNA-c1*. MRE start position within annotated *lncRNA-c1* transcript (TCONS_00034281) is indicated inside parenthesis. (E) Fold change in expression, of *GFP*, *GFP-lncRNA-c1* and *GFP-lncRNA-c1-MREΔ* (x-axis) in miRNA depleted cells transfected with negative control (NC) relative to miRNA depleted cells transfected with miRNA mimics (miRNA) (y-axis). Transcript expression was first normalized by the amount of *Act-β* and *PolII* and next by the total amount of transfected vectors per cell estimated based on the levels of relative *Neomycin* expression. Each point corresponds to the results of one independent biological replicate. Lines connecting data-points represent pairing of the three independent replicates. (F) Relative expression of *GFP-lncRNA-c1* and *GFP-lncRNA-c1-MREΔ* (x-axis) in wild-type mESC (WT). Transcript expression was first normalized by the amount of *Act-β* and *PolII* and next by the total amount of transfected vectors per cell estimated based on the levels of relative *Neomycin* expression. Each point corresponds to the results of one independent biological replicate. Statistics: *-p<0.05, **-p<0.01 and ** *-p<0.001 two-tailed paired t-test.

