## Supplementary figures and images for "Translation is required for miRNA-dependent decay of endogenous transcripts"

### Supplementary Figure S1

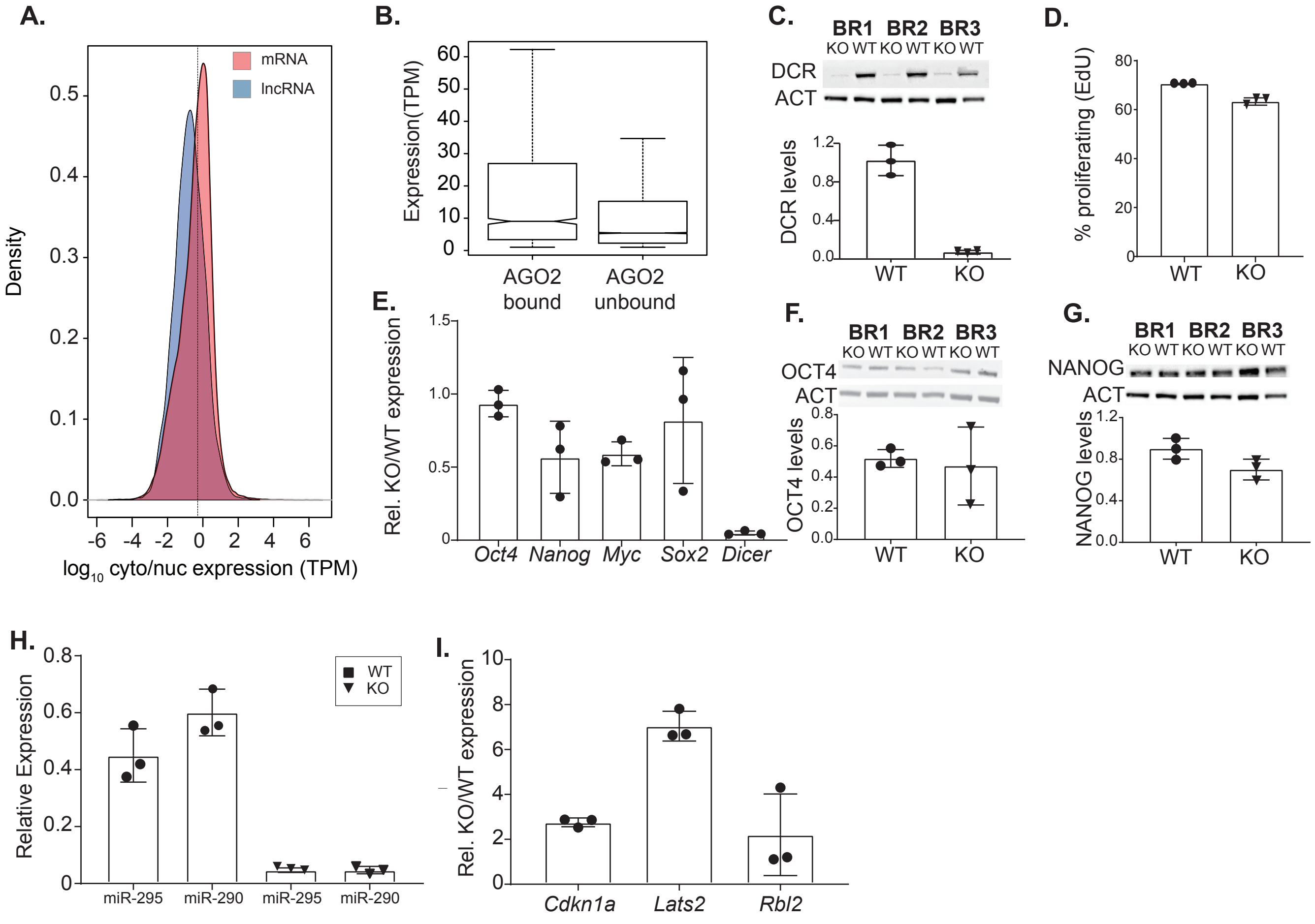

### Supplementary Figure S2

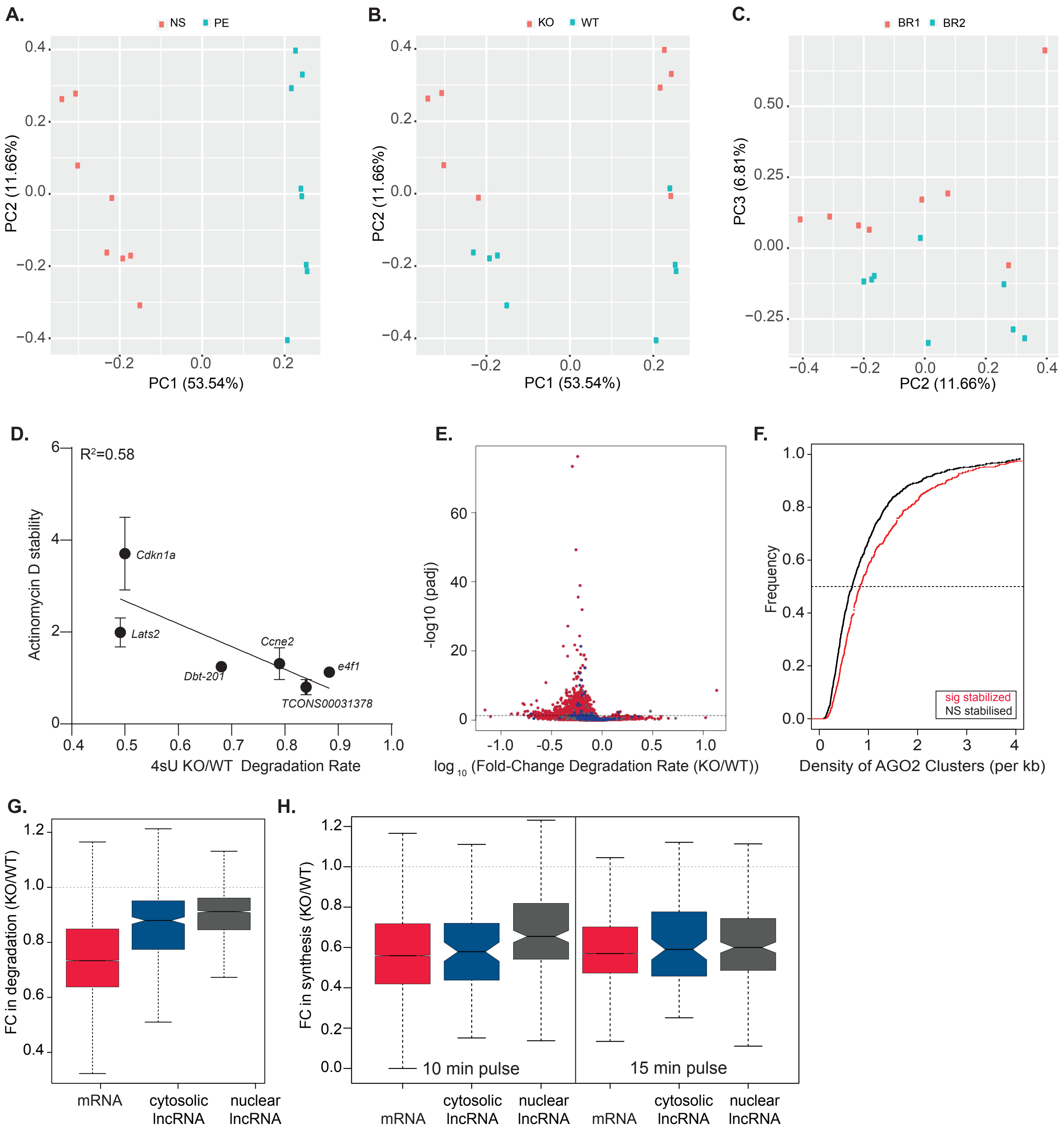

### Supplementary Figure S3

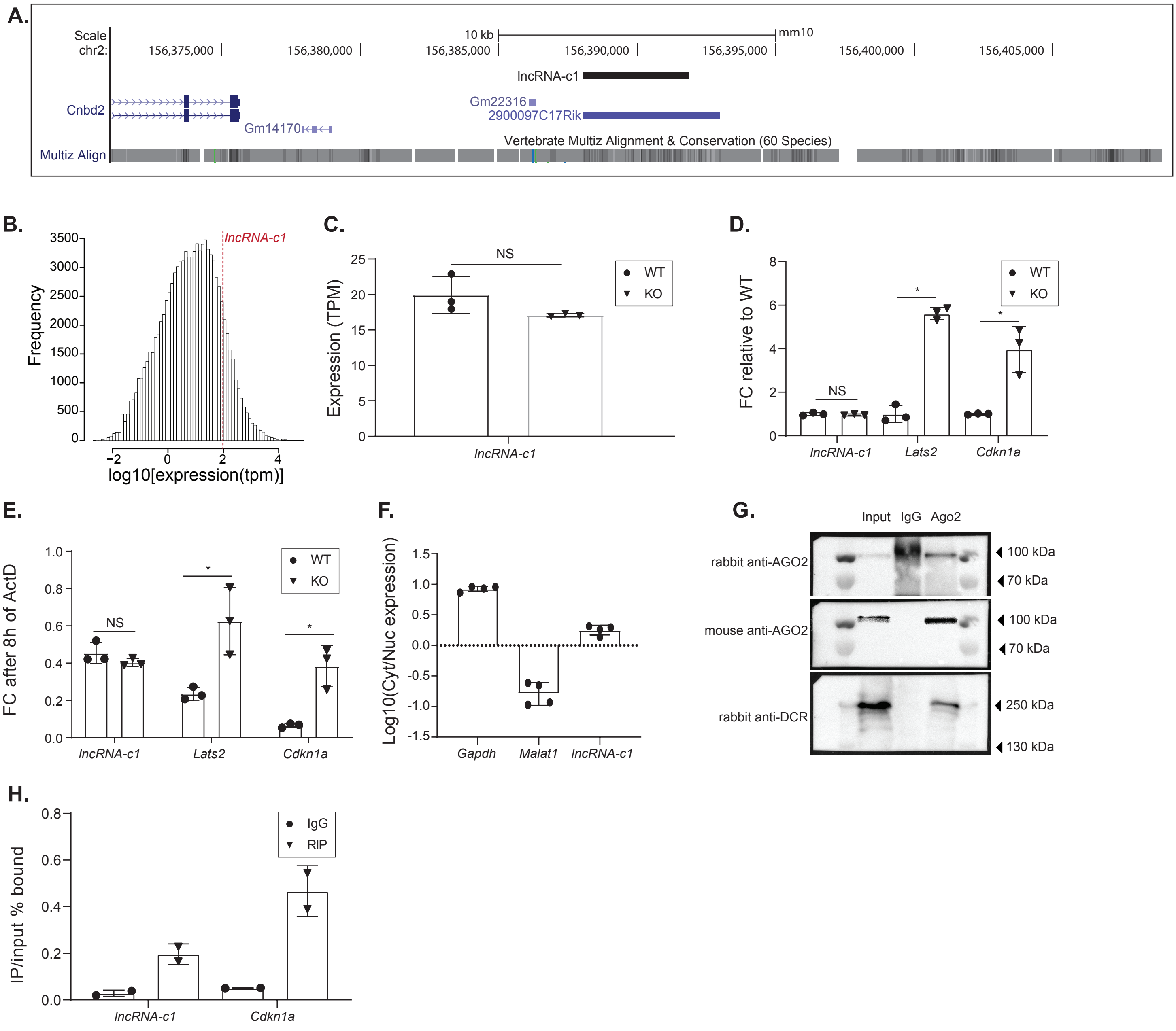
