## Supplementary Figure S4 for "Translation is required for miRNA-dependent decay of endogenous transcripts"

**A.**

T7 promoter / XhoI

*Cdkn1a* **TAATACGACTCACTATAG**GGAGACCCAAGCTGGCTAGCGTTTAAACGGGCCCTCTAGA**CTCGAG**ATGTCCAATCCTGGTGATGTC

*Cdkn1a∆ATG* **TAATACGACTCACTATAG**GGAGACCCAAGCTGGCTAGCGTTTAAACGGGCCCTCTAGA**CTCGAG**-TGTCCAATCCTGGTGATGTC

**B.**

```
MSNPGDVRPVPHRSKVCRCLFGPVDSEQLRRDCDALMAGCLQEARERWNFDFVTETPLEGNFVWERV
RSLGLPKVYLSPGSRSRDDLGGDKRPSTSSALLQGPAPEDHVALSLSCTLVSERPEDSPGGPGTSQG
RKRRQTSLTDFYHSKRRLVFCKRKP-SAHGSPALFCCGSGGLFPIFGLSPHSVCLNYYLCFNLNVSC
IYAACPLPVSKLKVI-KKNKTKKNQNKPKLVGR-GP-CGGFLLCRLLLFKPLPTQALCFLYRRNSPT
DIIPSASVSPNPPPPIPCLVPCHFLPGGDPQT-IALWKNE-DFGVSLSPLRPARMTVKQSQPRTGMA
VRTQP-YPDS-HCSDL-RQEWSPLWIPFATPGEPTSPVGLCQLPLYFGGLIW-SAALFPHPILPLLQ
VGRRHI-ALAPQLSGLEGNVYAGYTKWDSLVLP-AAPVATPCIVGLGWVLGGETGLPEHSMVCGGGG
GLIWDGDPSWGSQ-LLPFLSSSCTRSQAKMVSWGLRELTGH-AMADPFSVLNTVGVKALSGSDSSPK
HPCFCNILVWTVYP-PAPQEHVLWLPPCLHSDCKRLTRRDSTLHCPESSHPTPKLVLNKYFSMIEFH
HTGLVDPSSV
```

**C.**

```
CPILVMSDLFRTGAKCAVVSSVPWTVSSCAVIAMRSWRAVSRRPENGGTLTSSRRRRWRATSSGSAF
GA-GCPRST-ALGPAAVTTWEGTRGPVLPLPCCRGQLRRTTWPCRCLALWCLSGLKIPRVGPEHLRA
ENGGRPA-QISITPSADWSSARENPEVPTGAPPSSAVGQEASSPSSALALTLCVLIIICVLI-TSPV
YTLPALSQSPNLKLFKKRTKQKKTKTNLN--DGRALSVGDFYYVDYYYLSPSQPKLCVSYTGGTVLL
ISSHLHPFHPTPLPPFPAWFLATSYLGVILRPE-HFGKMSRTLGSPCHL-GQLG-Q-SSHSLEQGWQ
LGLNRNIPTLDIAQTCEDRSGPHSGSPLPLLGSPPLLWVSASCPSILEG-SGDLLLFSPTPYFPFCR
SAGGISRHLPHSSVDWKGMYMQGTLSGIPWSYLRQLQWQPPALWV-GGSLVVRQASQSILWCVVVGV
GLSGMGTPVGVLSDFSHFLVAVVQGARPRWCLGG-GSSQDTEQWLILSQC-IPWVSKHLVGLTPAPN
IPVSVTSWSGLSTLSPHPKNMYCGSLPVSTQIVSVSREGTAPCIVPSPHTRPQSWCSINTSR-LNST
TLD-XXRARYQ
```

**D.**

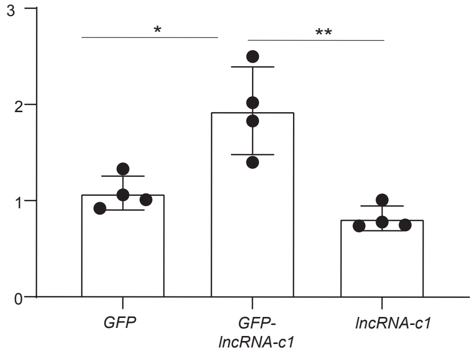


**E.**

```
      miR-295       3' agacucaaaaguaguAGCACUUu 5'
                                      ||||||||
lncRNA-c1 (1519)    5' cagauguucccaguaAGCACUUu 3'
lncRNA-c1 (2707)    5' ccuuuaaauguaucaAGCACUUu 3'

      miR-290       3' gggcuuaaaacuaggcGGCACUUu 5'
                                       |||||||
lncRNA-c1 (3314)    5' cauuuaacauacguugGGCACUUc 3'
```

**F.**

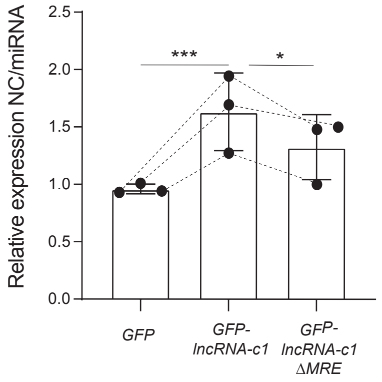


**G.**

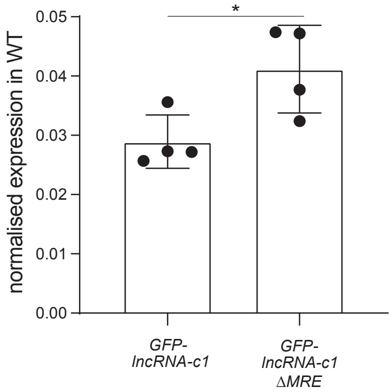
