## Supplementary File 1 for "Translation is required for miRNA-dependent decay of endogenous transcripts"

Supplementary File 1- Uncropped blots for NANOG and DICER detection

**Ponceau Rouge**

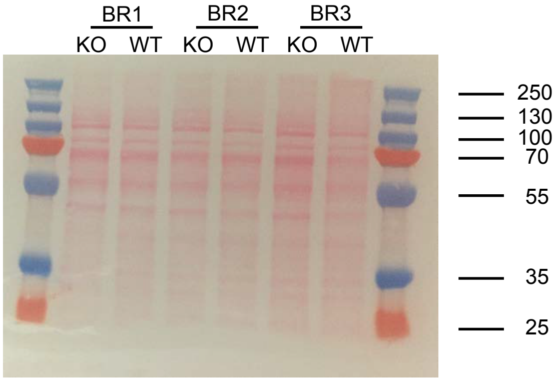


**Membrane was cut at 100 kDa to allow probing of DCR (214 kDa) and NANOG (35 kD)**

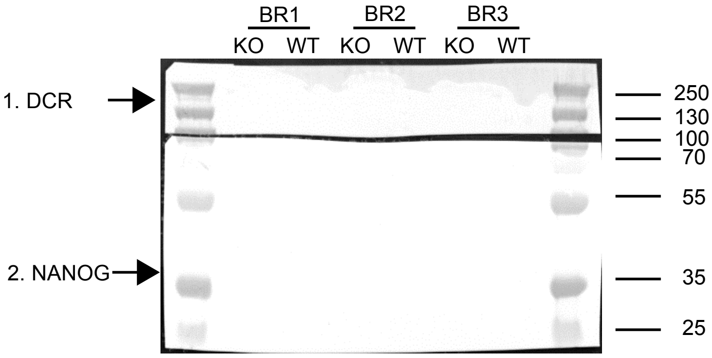


**Uncropped blot**

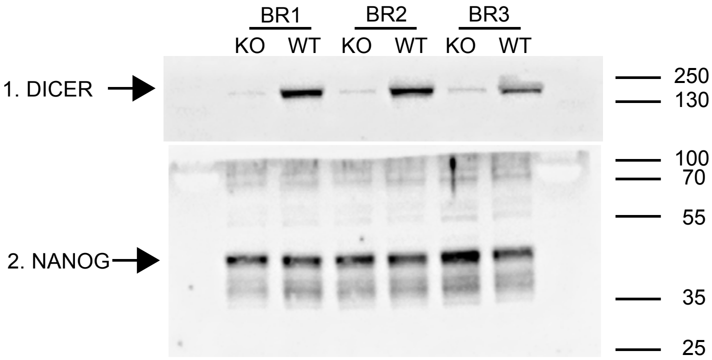


**Lower part of the membrane (2. NANOG) was stripped and probed for ACTIN-β (42 kDa)**

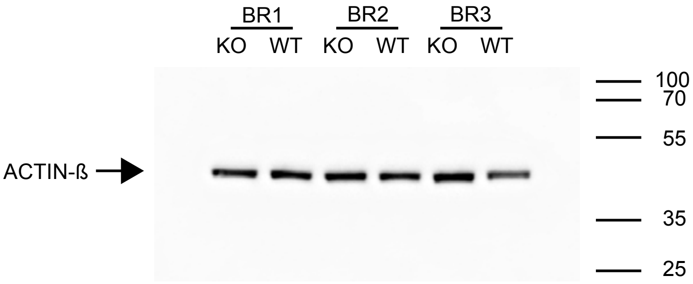
