## Supplementary File 2 for "Translation is required for miRNA-dependent decay of endogenous transcripts"

### Supplementary File 2- Uncropped OCT4 blots

**Ponceau Rouge**

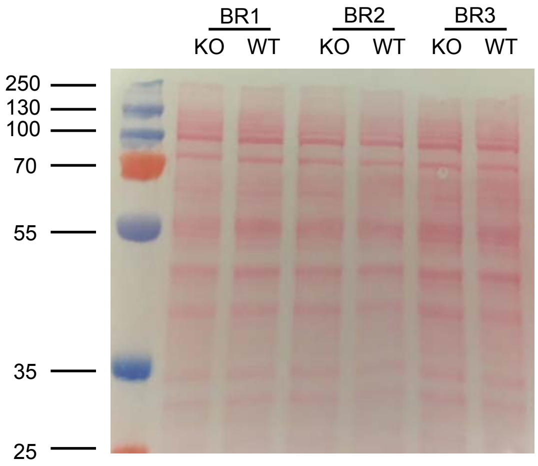


**Membrane was cut at 70 kDa- OCT4 (45 kDa) and ACTIN-β (42 kDa)**

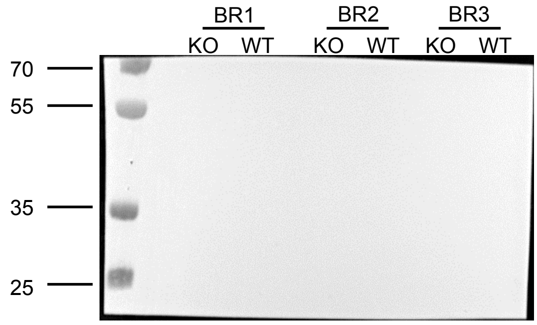


**Uncropped OCT4 blot**

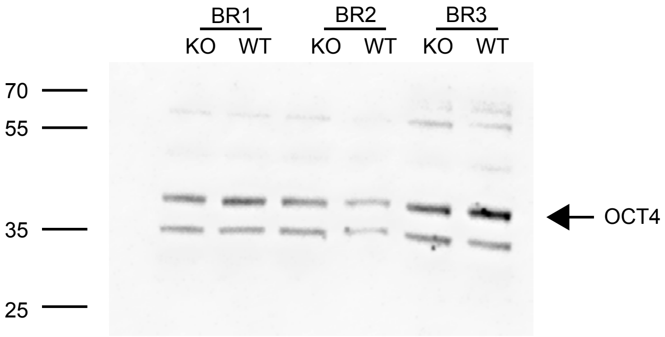


**Membrane was striped and probed for ACTIN-β. Uncropped blot below.**

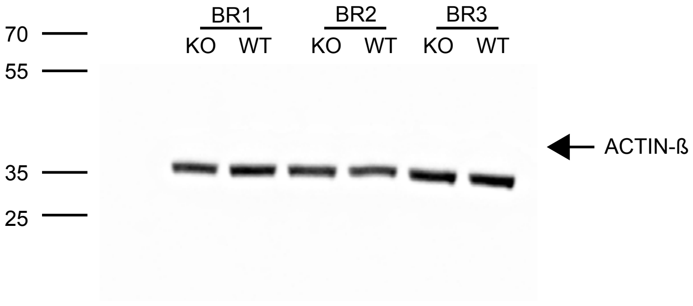
